# DANCE: A Deep Learning Library and Benchmark Platform for Single-Cell Analysis

**DOI:** 10.1101/2022.10.19.512741

**Authors:** Jiayuan Ding, Hongzhi Wen, Wenzhuo Tang, Renming Liu, Zhaoheng Li, Julian Venegas, Runze Su, Dylan Molho, Wei Jin, Wangyang Zuo, Yixin Wang, Robert Yang, Yuying Xie, Jiliang Tang

## Abstract

In the realm of single-cell analysis, computational approaches have brought an increasing number of fantastic prospects for innovation and invention. Meanwhile, it also presents enormous hurdles to reproducing the results of these models due to their diversity and complexity. In addition, the lack of gold-standard benchmark datasets, metrics, and implementations prevents systematic evaluations and fair comparisons of available methods. Thus, we introduce the DANCE platform, the first standard, generic, and extensible benchmark platform for accessing and evaluating computational methods across the spectrum of benchmark datasets for numerous single-cell analysis tasks. Currently, DANCE supports 3 modules and 8 popular tasks with 32 state-of-art methods on 21 benchmark datasets. People can easily reproduce the results of supported algorithms across major benchmark datasets via minimal efforts (e.g., only one command line). In addition, DANCE provides an ecosystem of deep learning architectures and tools for researchers to develop their own models conveniently. The goal of DANCE is to accelerate the development of deep learning models with complete validation and facilitate the overall advancement of single-cell analysis research. DANCE is an open-source python package that welcomes all kinds of contributions. All resources are integrated and available at https://omicsml.ai/.

## 1 Introduction

Single-cell profiling technology has undergone rapid development in recent years, spanning from single modality profiling (RNA, protein, and open chromatin) [19, 41, 62, 67, 74, 95, 102, 117, 129], multimodal profiling [14, 24, 54, 91, 143] to spatial transcriptomics [7,22,28,89,96,130,138,142]. The fast revolution in this field has encouraged an explosion in the number of computational methods, especially machine learning-based methods. However, the diversity and complexity of current methods make it difficult for researchers to reproduce the results as shown in the original papers. The major challenges include no publicly available code-base, hyperparameter tuning, and differences between programming languages. Furthermore, a systematic benchmarking procedure is necessary to comprehensively evaluate methods since the majority of existing works have only reported their performance on limited datasets and comparison with insufficient methods. Therefore, a generic and extensible benchmark platform with comprehensive benchmark datasets and metric evaluation is highly desired to easily reproduce any algorithm other than state-of-art methods under different tasks across popular benchmark datasets via minimal efforts (e.g., only one command line). Considering deep learning methods like Graph Neural Networks [26,118,121,140,143] have shown promising performance in single-cell analysis, but the customized interfaces of such tools are largely missing in the existing packages. Those motivate the development of our DANCE system that not only acts as a benchmark platform, but also provides customized deep learning infrastructure interfaces to help researchers conveniently develop their models.

In this work, we present DANCE as a deep learning library and benchmark to facilitate research and development for single cell analysis. DANCE provides an end-to-end toolkit to facilitate single cell analysis algorithm development and fair performance comparison on different benchmark datasets. DANCE currently supports 3 modules, 8 tasks, 32 models and 21 datasets. Table 1 summarizes the key differences between DANCE and existing single-cell libraries and toolkits. The highlights of DANCE are summarized as follows:

– **Comprehensive Module Coverage:** Squidpy [101] proposes an efficient and scalable infrastructure only for spatial omics analysis. DeepCell [136] forms a deep learning library for single-cell analysis but only biological images are covered. The library specializes in models for cell segmentation and cell tracking. Even though the popular Scanpy [144] provides a powerful tool for single-cell analysis spanning all modules, it focuses on the field of data preprocessing instead of modeling. Similarly, even though Seurat [54] touches on all three modules, its R language-based interface restricts its applicability for the development of deep learning methods due to limited R interface support within the deep learning community. Instead, DANCE supports all types of data preprocessing and modeling across all modules including single modality, multimodality and spatial transcriptomics.
– **Deep Learning Infrastructure:** With the great increase in the number of single cells, classical methods [15, 68] cannot effectively enjoy the benefit from big single-cell data, while deep learning has been proven to be effective. Furthermore, deep learning techniques are also good at handling high dimensional data, which is common for single-cell data. Unfortunately, the backend framework of the well-known Seurat is R, which limits its potential in the deep learning community due to restricted R interface support in the deep learning community. Scanpy only contains classical methodologies for downstream tasks. Recently scvi-tools [43] presents a Python library for deep probabilistic analysis of single-cell omics data. With 12 models, scvi-tools offers standardized access to 9 tasks. scvi-tools includes some deep learning methods but lacks the recent Graph Neural Networks (GNNs) based methods. In terms of models, scvi-tools selects baselines with a concentration on statistical models according to their supporting data protocol. As a comparison, DANCE is a comprehensive deep learning library of single-cell analysis. Popular deep learning infrastructures like Autoencoders [116] and GNNs are supported and applicable for all modules.
– **Standardized Benchmarks:** To the best of our knowledge, DANCE is the first comprehensive benchmark platform covering all modules in single-cell analysis. A few unique features have been developed to achieve this goal. We first collect task specific standard benchmark datasets and provide easy access to them by simply changing the parameter setting. Under each task, representative classical and deep learning algorithms are implemented as baselines. Those baselines are further fine-tuned on all collected benchmark datasets to reproduce similar or even better performance compared to original papers. To easily reproduce the results of our finetuned models, end users only need to run one command line where we wrap all super-parameters in advance to obtain reported performance.

**Table 1:**
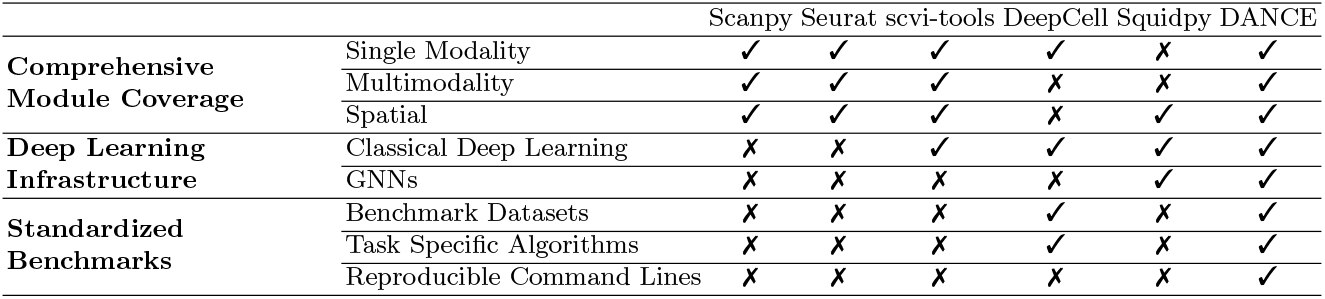
Comparison between DANCE and other popular single-cell libraries and toolkits.

One of the highlights of DANCE is the reproducibility of models. The diverse programming languages and backend frameworks of existing methods make systematic benchmark evaluation challenging for fair performance comparison. In such case, we implement all models in a unified development environment based on python language using Pytorch [105], Deep Graph Library (DGL) [141] and PyTorch Geometric (PyG) [40] as backbone frameworks. In addition, we formulate all baselines into a generic fit-predict-score paradigm. From the reproducibility perspective, for each task, every implemented algorithm is fine-tuned on all collected standard benchmarks via grid search to get the best model, and the corresponding super-parameters are saved into only one command line for user’s reproducibility. We also provide one example for each model as a reference.

## 2 Related Work

### 2.1 Single-Cell Analysis

#### Single Modality

Next-generation sequencing allows for high-throughput transcriptome analysis. While bulk RNA sequencing data focuses on average gene expression profiles [76, 100, 110], single-cell RNA-sequencing (scRNA-seq) data quantifies gene expression at the cell level, offering unprecedented advantages to understanding biological mechanisms. In scRNA-seq experiments, single cells need to be isolated and captured; then, the captured cells go through lysis [66, 76, 100]. Reverse transcription is then applied to lysed cells for mRNA selection and cDNA synthesis, which is a key step that determines experiment sensitivity and the level of technical noise due to sampling noise, which is often assumed to have a Poisson distribution [58]. The rapid development of sequencing technologies has enabled highly scalable experiments where a massive number of cells could be processed simultaneously through microwell-based methods such as CytoSeq [58], microfluidics-based methods like Fluidigm C1 HT, or droplet-based methods like Drop-seq [86] and Chromium 10X [155]. In particular, the development of unique molecular identifiers (UMIs) improves the quality of scRNA-seq by barcoding each mRNA molecule within a cell individually during reverse transcription and thus mitigates PCR amplification bias. In addition to scRNA-sequencing, other types of single-cell sequencing assays also emerged recently. Two common branches are sequencing for surface protein such as CITE-seq [124] and sequencing for chromatin accessibility as represented by scATAC-seq [70].

Single-cell sequencing technology enables the simultaneous screening of thousands of genes in a massive number of cells and yields valuable insights in cell heterogeneity and functions. For instance, it has shed light on new understandings in oncological studies [72,88,145], cell characterizations [27,48,49,112,133,134,137], immune heterogeneity [44, 103, 154], and cell differentiation [12, 69, 97, 115]. In order to answer important biological questions, new algorithms are required that are specially devised for single-cell sequencing data. Unlike bulk sequencing, single-cell sequencing technologies profile thousands or even millions of cells, resulting in high-dimensional datasets. Furthermore, single-cell sequencing data suffers from high sparsity [71], i.e, a large proportion of genes with zero reads in the count matrix, a phenomenon known as dropout [8, 63, 66, 113]. Thus, new algorithms are needed to efficiently analyze single-cell sequencing data in order to tackle tasks such as data imputation, cell clustering, cell typing, and cell trajectory inferences.

#### Multimodality

There have been exploding experiment technologies that obtain high-resolution features of single-cell, and many of them devoted to adapting assays of different omics to single-cell resolution (e.g., scDNA-seq [87], scATAC-seq [70], REAP-seq [108], etc)Recently, advances in single-cell technology have led to simultaneous assays of multi-omics in a single cell. For example, cellular indexing of transcriptome and epitopes by sequencing (CITE-seq) [124] simultaneously profiles mRNA gene expression and surface protein abundance; sci-CAR [16] and SNARE-seq [23] jointly measure mRNA gene expression and chromatin accessibility. Integrated analysis of multi-omics single-cell data has brought many important applications and achievements, such as revealing new cell populations [54] or regulatory networks [37, 59, 82, 153]. Current multi-omics analysis can be roughly categorized into two types, i.e., the joint analysis of data from different single-modal sources, and the analysis of data from multi-modal assays. Both of them can provide new insights into cell states. The most important problem in multimodal data analysis is how to integrate multimodal data to provide more accurate and in-depth cellular representation. To this end, integration algorithms can also be considered in two categories. The first category of methods [18, 37, 59, 61, 77, 122, 152, 153] focus on alignment between datasets, while the second category of methods [92, 143, 146, 149, 158] dedicate to capture cellular characterization from multimodality.

Although various successful downstream applications have demonstrated the effectiveness of multimodal data, it is still difficult to directly evaluate and compare the performance of different integration methods. One common way to evaluate those methods is to calculate Normalized Mutual Information (NMI) between the predicted cluster labels and predefined cell type labels. A more reasonable way to benchmark multimodal integration, as suggested in a recent work [80], is to leverage multi-omics aligned data to provide ground truth for multimodal integration, where two modalities are simultaneously measured in each cell (e.g. CITE-seq [124]). Three key tasks have been defined(i.e., modality prediction, modality matching, and joint embedding) to comprehensively evaluate the power of various integration methods. Modality prediction is to predict one modality from another. In modality matching, we aim to identify observations of the same cells among different modalities, while the ground truth correspondence is given by the dataset. Joint embedding is to directly evaluate the integrated embedding by comprehensive metrics based on biological states preservation and batch effects removal.

#### Spatial Transcriptomics

While single-cell sequencing technologies are able to isolate cells from heterogeneous cell populations, they lose sight of the original microenvironments in the process. Spatial transcriptomics technologies are a more recent development in transcriptomics profiling. They are able to spatially resolve transcriptomics profiles. This allows researchers to further investigate and understand the spatial context of cells, and cell clusters [13]. However, the spatially resolved profiling regions for most technologies are not at single-cell resolution, which motivates the problem of spatial (cell-type) deconvolution and segmentation.

Spatially resolved RNA profiling technologies can be roughly categorized as profiler-based and imager-based technologies [55]. Profiler-based technologies use Next-Generation Sequencing (NGS) readouts, and include Spatial Transcriptomics (10x Visium platform) [126], Slide-seqV2 [123], and Digital Spatial Profiling (Nanonstring GeoMx platform) [90]. The profiler-based technologies offer high-plex data, though not at single-cell resolution. Imager-based technologies apply a sequence of cyclic In situ hybridization (ISH) and imaging. Recent methods include MERFISH (Visgen MERSCOPE platform) [93], seqFISH+ [39], and the Spatial Molecular Imager (Nanostring CosMx platform) [55]. Imager-based technologies offer single-cell resolution (even sub-cellular), though usually not as high-plex as the profiler-based methods [55].

### 2.2 Graph Representation Learning

Graph representation learning has attracted increasing attention [84], since graph data are ubiquitous in the real world, e.g. social networks [98], knowledge graphs [60] and biological networks [106]. However, graph data has a more complex topology than text or image data with regular structures, causing it difficult to analyze. To facilitate downstream analysis, a natural idea is to denote each node with a vector while encoding intrinsic graph properties. This kind of methods is called graph embedding [84], including random-walk-based methods [47, 107], matrix-factorization-based methods [4, 10], and deep learning methods [147].

Random walk approaches (e.g., deepwalk [107], node2vec [47]) start with sampling the neighborhoods of nodes rather than directly using the global information of the entire graph. This allows the model to capture higher-order relationships between nodes, but they also lose some global structural information. Matrix factorization methods (e.g., graph laplacian eigenmaps [10], graph factorization [4]) have a solid mathematical foundation, but they are not scalable to large graphs, due to the high space complexity of proximity matrix construction and computational complexity of eigen decomposition. Moreover, most factorization methods only conserve the first-order proximity [20]. Deep learning models, also known as graph neural networks (e.g., GCN [64], GraphSAGE [50]), are generally achieving state-of-the-art performance in various applications. However, they are mathematically complex and thus lack interpretability. To be concrete, graph neural works (GNNs) iteratively propagate and transform node features to obtain node embeddings [46]. Therefore the node embeddings encode high-order topological structure information. In addition, GNNs provide denoising effects by smoothing graph signals through filtering eigenvalues of the graph Laplacian [83]. Because of these advantages, graph representation learning can be very effective for single-cell analysis. For instance, it can facilitate cell-cell interaction detection [73] and gene-network inference [59].

### 2.3 Deep Learning for Single Cell Analysis

In contrast to the bulk sampling techniques, single-cell sequencing techniques can produce millions of samples in a single experiment, which far exceeds the sample size of previous datasets. Along with this is the emergence of a large number of new research topics, mainly focusing on understanding the association between gene expressions and cellular behaviors. These changes have led researchers to focus on machine learning algorithms, which make good use of large training sets to optimize a given objective function. Particularly, deep learning methods consistently show outstanding performance in numerous machine learning applications [34, 111]. As a result, deep learning models are now widely used in single-cell analysis and have greatly aided the development of immunology, oncology, pharmacology, and many other disciplines [21, 45, 135].

The enormous potential of single-cell data comes with great complexity caused by various underlying biological and technical factors. To address this issue, several pre-processing steps are developed into the analysis pipeline. In the early stages, quality control and normalization are carefully designed. After that, complex machine learning tasks are introduced, e.g., batch effect correction, data imputation, and dimension reduction. Special types of single-cell data may require further processing, such as multimodal data integration for multi-omics and cell deconvolution for spatial transcriptomics. All these tasks can be facilitated by deep learning methods [6, 79, 114, 139]. Meanwhile, Deep learning methods [26,32,36,42,81,99,128,131,132,148,151] consistently outperform other classical machine learning techniques in downstream tasks, including clustering, cell type annotation, disease prediction, gene network inference, and trajectory analysis.

## 3 An Overview of DANCE

### 3.1 Environment Requirements and Setup

DANCE works on python ≥ 3.8 and Pytorch ≥ 1.11.0. All dependencies are listed in Appendix A. After cloning this repository, run setup.py to install DANCE into the local python environment or install it directly from pip install as below:

~~~
pip install pydance
~~~

### 3.2 The Architecture Design

Figure 1 provides an overall design of the architecture of the DANCE package. The DANCE package consists of two key components: lower-level infrastructure and upper-level task development.

**Fig. 1:**
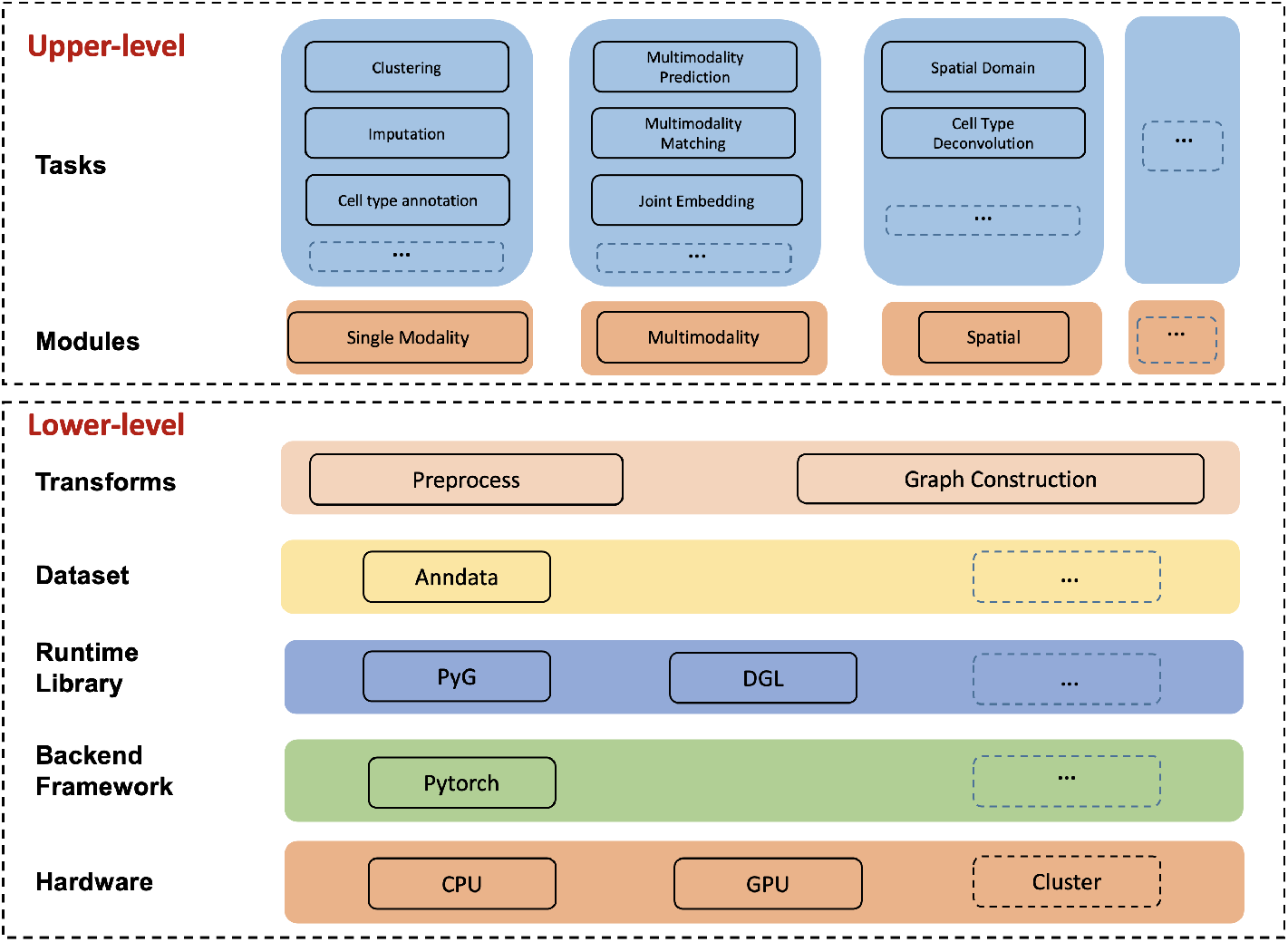
The architecture of DANCE package.

#### Lower-level Infrastructure

From the hardware perspective, CPU running is supported for all methods developed in DANCE. In addition, for deep learning based methods, we also support GPU running to accelerate the training process, especially for large-scale datasets. In the future, cluster running for deep learning methods would be also developed to support model training across multiple GPUs. The backbone framework in DANCE is Pytorch [105], which is used for high-performance deep learning model development. To support various methods for deep learning on graphs and other irregular structures, we take both Deep Graph Library (DGL) [141] and PyTorch Geometric (PyG) [40] as graph engines in DANCE. Various types of preprocessing functionalities are provided in the Transforms folder to process data before model training. For methods based on Graph Neural Networks (GNNs), we also support distinct ways of graph construction to convert cell-gene data like RNA sequencing (RNA-seq) to cell-cell, cell-gene and gene-gene graphs. What’s more, spatial coordinates and image features of single cells can be also extracted to help construct graphs for spatial transcriptomics. Those lower-level interfaces are helpful for developers to build their models on downstream tasks without building “wheels” from scratch.

#### Upper-level Task Development

Based on the infrastructure described above, individual modules and tasks can be further defined and developed. Currently, we support tasks under single modality profiling, multimodal profiling, and spatial transcriptomics modules, which correspond to three stages of single-cell technology development. Under each module, classic tasks are covered, and representative methods are implemented through the evaluation on several standard benchmarks. Note that upper-level task development is highly flexible and extensible. This indicates that users can readily extend their new modules, tasks, models, and datasets into the existing repository of DANCE.

### 3.3 A Pipeline of DANCE

#### Data Loading

For each task, we have a generic interface to load datasets. All datasets supported by DANCE are cached on the cloud. Users don’t have to download their interested datasets manually. They just need to specify an individual dataset when calling the data loader interface. For example, we can run graph-sc model on **10X PBMC** dataset for the clustering task using the following command line:

~~~
python graphsc .py -- dataset =‘10 X_PBMC’
~~~

#### Data Processing

After data loading, a collection of data processing methods is provided before model training. They are divided into two parts: preprocessing and graph construction.

– **Preprocessing:** We provide rich preprocessing functions such as normalization, dimension reduction, gene filtering and so on. Take graph-sc model as an example, we filter out the rarely expressed genes and normalize the remaining to obtain the same total count for each cell. Then only the highly expressed genes (top 3000 by default) are kept for clustering [26].
– **Graph Construction:** This is required for GNNs based method. Before model training, we have to convert data to graphs in preparation for graph operations. DANCE provides a variety of ways of graph construction. In graph-sc implementation, we construct a weighted heterogeneous cell-to-gene graph, where the types of nodes can be cell and gene nodes. The weight of each edge between each cell node and its corresponding gene node is determined by gene counts, and there is no edge linked between any pairs of cell or gene nodes.

#### Model Training and Evaluation

All models in DANCE have generic interfaces for model training and evaluation. The unified interface for model training is **model.fit()** while that for model prediction is **model.predict()**, which returns the predictions of test data. Furthermore, **model.score()** acts as a generic interface to evaluate how well each model is. The metric of the score function depends on each task. Take graph-sc for an example, after fitting the model with chosen hyperparameters, we can access the performance of graph-sc by calling the score function, which will return ARI and NMI scores to indicate the quality of the clusters.

## 4 DANCE Benchmark: Modules, Tasks, Models, and Datasets

As shown in Figure 2, DANCE is capable of supporting modules of single modality, multimodality and spatial transcriptomics. Under each module, we benchmark several tasks with popular models across standard datasets. Here, we take the task of Clustering in the module of single modality as an example. Various types of methods are implemented including Graph Neural Networks based methods including graph-sc [26], scTAG [151] and scDSC [42] and AutoEncoders based methods including scDeepCluster [131] and scDCC [132]. To ensure a systematic evaluation and fair performance comparison of different models, several standard benchmark datasets such as 10X PBMC 4K [156], Mouse Bladder Cells [51], Worm Neuron Cells [17] and Mouse Embryonic Stem CellS [65] for the task are collected for evaluation. Currently, there are 3 modules, 8 tasks, 32 models and 21 datasets supported by DANCE. Please refer to Appendix D and Appendix C for more details about supported models and datasets, respectively.

**Fig. 2:**
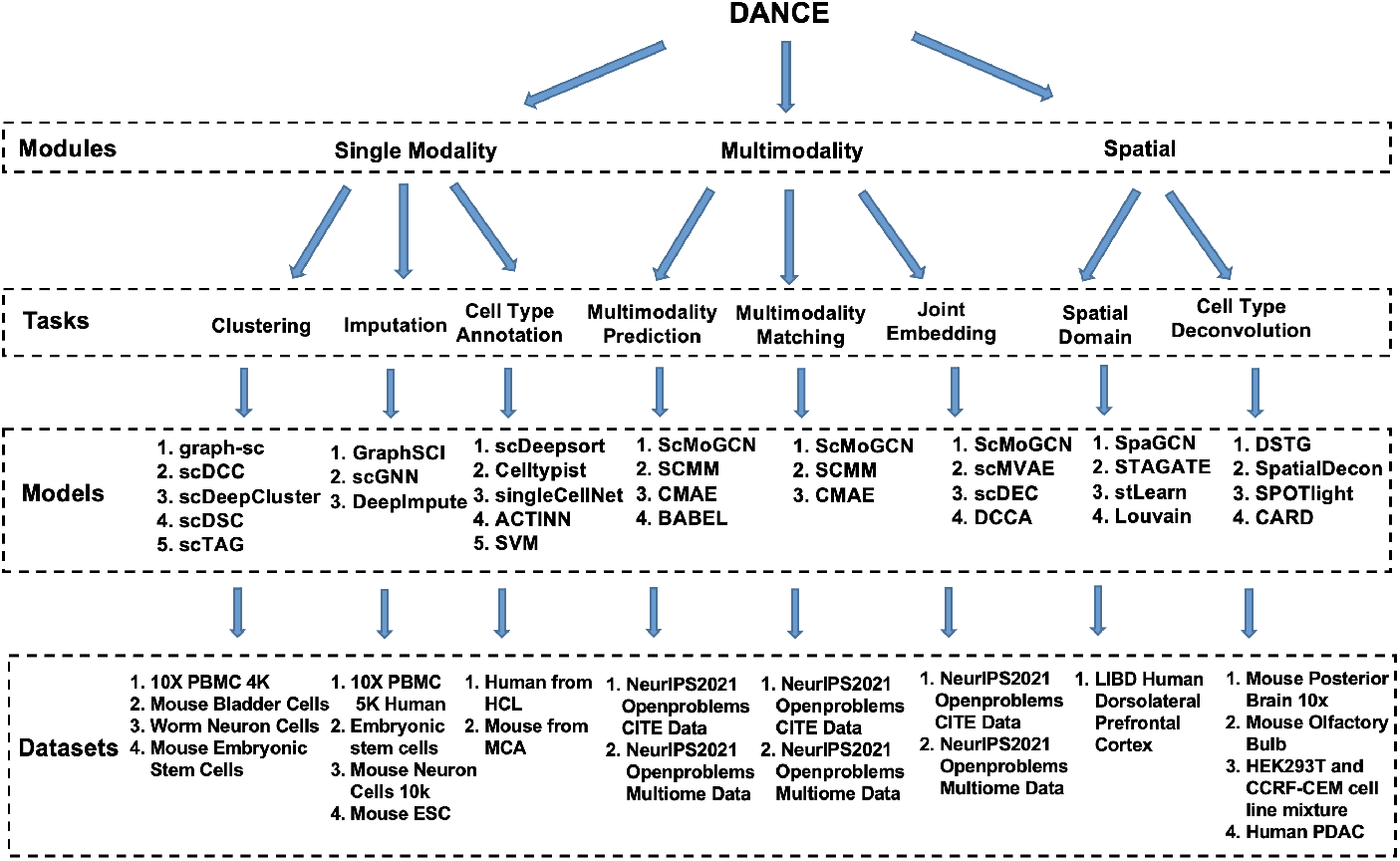
A summary of modules, tasks, models and datasets supported by the DANCE package.

## 5 Benchmarking and Reproduction

One of the highlights of DANCE is the reproducibility and potential benchmarking. It is always challenging but helpful to reproduce the performance of the published works in single-cell analysis since the implementations are based on different programming languages and different backend frameworks. To tackle this challenge, we implement all models based on Pytorch, PyG and DGL, and put all baselines into the fit-predict-score structure. For every task, we choose several benchmark datasets and tune all baselines on every dataset. When implementing baselines, we refer to the implementation details in the original GitHub repositories and the corresponding papers. For models without suggested parameters, we perform grid search and random search to obtain the best parameters. All the parameter settings can be found via https://github.com/OmicsML/dance/tree/main/examples, where we keep command line information for reproduction at the end of every example file.

To illustrate reproducibility and benchmarking in DANCE, we delve deeper into the cell type annotation task. Note that all reproductions are accessible on GitHub: https://github.com/OmicsML/dance. Currently, DANCE supports five models for this task. It includes scDeepsort [118] as a GNN-based method. ACTINN [81] and singleCellNet [128] are representative deep learning methods. We also cover support vector machine (SVM) and Celltypist [32] as traditional machine learning baselines. We select datasets on which all five models have reported results, i.e., MCA [52] for mouse and HCL [53] for human. As for the evaluation purpose, we pull three tissues from the MCA and pick one dataset from each tissue, i.e., Mouse Brain 2695, Mouse Spleen 1759 and Mouse Kidney 203.

Table 2 demonstrates the reproduced and reported results of the cell-type annotation task in terms of accuracy (ACC). For every table cell, on the left, we have the reproduced annotation ACC based on DANCE, and on the right, we show the reported ACC in the original works. As shown in Table 2, SVM and scDeepsort [118] get similar results as reported. Impressively, Celltypist [32] and ACTINN [81] bring out advancement under the DANCE implementation with an average absolute increase of 7% and 44% over three datasets, respectively. Note that only singleCellNet [128] produces lower ACC compared to the reported results. Among all supported tasks, DANCE achieves uniform benchmarking grounded in Pytorch, which requires less configuration and allows users to conduct analysis solely using Python. The structure of some models may be modified in the process of transformation from R or TensorFlow to Pytorch, which in consequence affects the reproduced results (e.g., singleCellNet).

**Table 2:** Reproduced and reported ACC for the cell type annotation task. *To meet the format requirements, we renormalize datasets before running the original implementation of Celltypist.

| Model | Mouse Brain 2695<br>(current / reported) | Mouse Spleen 1759<br>(current / reported) | Mouse Kidney 203<br>(current / reported) |
| --- | --- | --- | --- |
| scDeepSort | 0.363 / 0.363 | 0.965 / 0.965 | 0.901 / 0.911 |
| Celltypist* | 0.680 / 0.666 | 0.966 / 0.848 | 0.879 / 0.832 |
| singleCellNet | 0.693 / 0.803 | 0.975 / 0.975 | 0.795 / 0.842 |
| ACTINN | 0.860 / 0.778 | 0.516 / 0.236 | 0.829 / 0.798 |
| SVM | 0.683 / 0.683 | 0.056 / 0.049 | 0.704 / 0.695 |

## 6 Open-Source Contribution

DANCE is an open-source package, and everyone can contribute to this platform with extra tasks, models, and standard benchmark datasets by following the instructions below:

– **Codebase Structure Guidance:** Before contributing to DANCE, you need to understand the codebase structure in DANCE and make sure you are going to modify the correct files or put new files into the correct place. [dance](https://github.com/OmicsML/dance/tree/main/dance) is the root of the package. [datasets](https://github.com/OmicsML/dance/tree/main/dance/datasets) is the dataloader module, which contains task specific data loaders. [modules](https://github.com/OmicsML/dance/tree/main/dance/modules) is the model module, which covers all implemented algorithms. Each model is created as an individual file under the corresponding task folder. [transforms](https://github.com/OmicsML/dance/tree/main/dance/transforms) is the data processing module, which deals with data preprocessing in [preprocess](https://github.com/OmicsML/dance/blob/main/dance/transforms/preprocess.py) and graph construction in graph_construct. You are flexible to pick up any existing functionalities in those two files for your model development and welcome to contribute new ones if your desired ones are not provided. examples is the example reference module, which presents one example for one model.
– **Code Style Guidance:** The contributed codes are required to follow the standard python coding style. For more details, please refer to [Style Guide for Python Code](https://peps.python.org/pep-0008/).
– **Testing:** To ensure that your contributed code does not impact the normal operation of the existing codebase, you need to ensure that all tests pass before submitting. Please refer to the **Run Test** section in DANCE for how to run tests.
– **Reproducibility:** This is only required for contributed new models. An example file is necessary to present how to run your model on existing standard benchmarks. Furthermore, you need to place command lines at the end of the example file to obtain the best performance for reproducibility purposes. One command line corresponds to one standard benchmark dataset running of your contributed model.
– **Documentation:** Submitted code should be documented or commented on for easy readability purposes by users. Please refer to [Numpy Style Docstring Guide](https://numpydoc.readthedocs.io/en/latest/format.html) for more details.

## 7 Impacts and Future Directions

In face of reproducibility issues of computational models in the field of single-cell analysis, we believe the DANCE platform will bring meaningful contributions to the whole single-cell community. To be specific, end users don’t have to spend a lot of effort in implementing and tuning models. Instead, they only need to run the command line we provide to easily reproduce results from the original paper. In addition, with our implementation, the performance of some models is even better than that reported in the original paper; we also provide GPUs support for the accelerated training purpose for deep learning based models in our implementation. It is worth noting that our DANCE package is an open-source package where every developer can contribute to advancing this field.

Since the functionalities of preprocessing and graph construction in current DANCE are not consolidated. It would be enhanced in the future. DANCE would be released as a SaaS service, which means that users are not limited to their local computation and storage resources. The interactive interface will be also provided for end users to use. In such case, after users upload their datasets to the platform, they only need to click on some buttons to select the preprocessing functions and the interested tasks and models for running without coding skills requirement. The results would be visualized on the website instantly. The leaderboard can be further developed to evaluate the efficacy and generalizability of state-of-the-art computational methods. From the data perspective, the collected datasets enable us to integrate them into atlas reference database via the exploration of integrating different modalities or even tissues. Later on, this atlas would be released for public access to further research in the whole single-cell community. Last but not least, AutoML and model explainability will be supported which can further minimize the efforts and ML background requirements for DANCE users.

## A Environment Dependencies

For the dependencies of our first version of the DANCE package, please refer to Table 3 for more details.

**Table 3:** Dependencies of DANCE Package

| Dependency | Version |
| --- | --- |
| h5py | $\geq 3.7.0$ |
| leidenalg | $\geq 0.8.10$ |
| networkx | $\geq 2.8.5$ |
| numba | $\geq 0.56.0$ |
| opencv-python | $\geq 4.6.0.66$ |
| openpyxl | $\geq 3.0.10$ |
| psutil | $\geq 5.9.1$ |
| pyro-ppl | $\geq 1.8.1$ |
| python-igraph | $\geq 0.9.11$ |
| rdata | $\geq 0.8$ |
| scanpy | $\geq 1.9.1$ |
| scikit-learn | $\geq 1.1.2$ |
| scikit-misc | $\geq 0.1.4$ |
| scipy | $\geq 1.9.0$ |
| seaborn | $\geq 0.11.2$ |
| skorch | $\geq 0.11.0$ |
| statsmodels | $\geq 0.13.2$ |
| torch | $\geq 1.11.0$ |
| torchnmf | $\geq 0.3.4$ |
| torchvision | $\geq 0.12.0$ |
| tqdm | $\geq 4.64.0$ |

## B Performance Display

In this section, we show the performance comparison between our implementation and the original implementation for tasks as below:

– **Single Modality Module:** Imputation, Cell Type Annotation and Clustering.
– **Multi-modality Module:** Modality Prediction, Modality Matching and Joint Embedding
– **Spatial Module:** Spatial Domain and Cell Type Deconvolution.

If the original implementation is unavailable to the public or is based on non-python language, we re-implement them with generic python language in our DANCE package. If the original implementation exists, we transform them into the generic fit-predict-score structure. Note that not all methods are tested on all standard benchmark datasets. To bridge this gap, we test them on each standard benchmark dataset for system evaluation and fair comparison. In all performance comparison tables, we compare the performance of our re-implemented algorithms denoted as “current” with that of original ones denoted as “reported”. Note that N/A means that the performance is not available on this dataset.

**Table 4:**
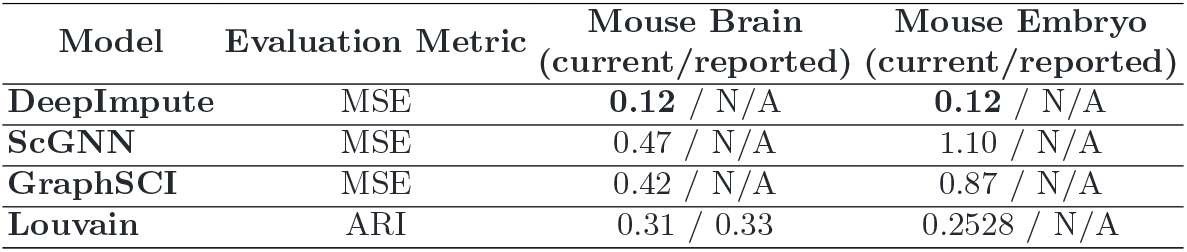
Imputation Task: performance comparison between our and original implementations in DANCE.

**Table 5:**
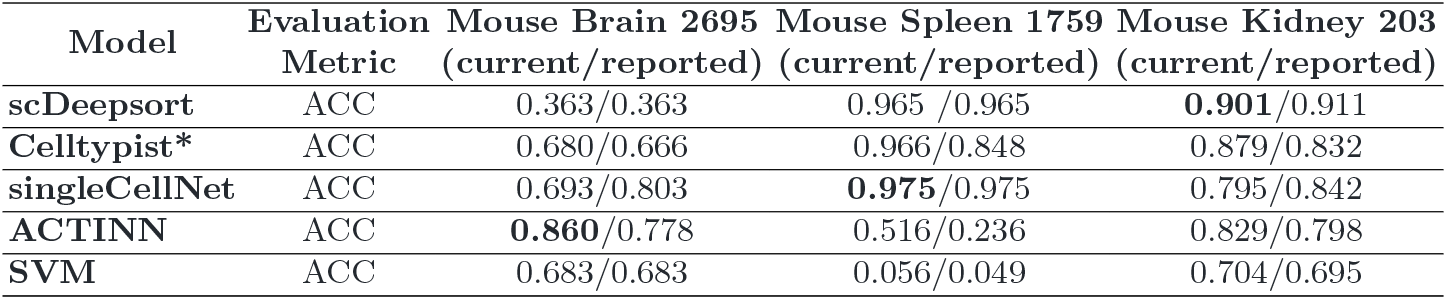
Cell Type Annotation: performance comparison between our and original implementations in DANCE. (Note: * Benchmark datasets have been renormalized when running the original implementation of Celltypist to meet its format requirements.)

| Model | Evaluation Metric | Mouse Brain 2695<br>(current/reported) | Mouse Spleen 1759<br>(current/reported) | Mouse Kidney 203<br>(current/reported) |
| --- | --- | --- | --- | --- |
| scDeepsort | ACC | 0.363/0.363 | 0.965 / 0.965 | **0.901**/0.911 |
| Celltypist* | ACC | 0.680/0.666 | 0.966/0.848 | 0.879/0.832 |
| singleCellNet | ACC | 0.693/0.803 | **0.975**/0.975 | 0.795/0.842 |
| ACTINN | ACC | **0.860**/0.778 | 0.516/0.236 | 0.829/0.798 |
| SVM | ACC | 0.683/0.683 | 0.056/0.049 | 0.704/0.695 |

**Table 6:** Clustering Task: performance comparison between our and original implementations in DANCE.

| Model | Evaluation Metric | 10x PBMC<br>(current/reported) | Mouse ES<br>(current/reported) | Worm Neuron<br>(current/reported) | Mouse Bladder<br>(current/reported) |
| --- | --- | --- | --- | --- | --- |
| graph-sc | ARI | 0.72 / 0.70 | 0.82 / 0.78 | **0.57** / 0.46 | **0.68** / 0.63 |
| scDCC | ARI | **0.82** / 0.81 | **0.98** / N/A | 0.51 / 0.58 | 0.60 / 0.66 |
| scDeepCluster | ARI | 0.81 / 0.78 | 0.98 / 0.97 | 0.51 / 0.52 | 0.56 / 0.58 |
| scDSC | ARI | 0.72 / 0.78 | 0.84 / N/A | 0.46 / 0.65 | 0.65 / 0.72 |
| scTAG | ARI | 0.75 / N/A | 0.96 / N/A | 0.53 / N/A | 0.60 / N/A |

**Table 7:** Modality Prediction Task: performance comparison between our and original implementations in DANCE.

| Model | Evaluation Metric | GEX2ADT<br>(current/reported) | ADT2GEX<br>(current/reported) | GEX2ATAC<br>(current/reported) | ATAC2GEX<br>(current/reported) |
| --- | --- | --- | --- | --- | --- |
| ScMoGCN | RMSE | **0.3885** / 0.3885 | **0.3242** / 0.3242 | **0.1778** / 0.1778 | **0.2315** / 0.2315 |
| SCMM | RMSE | 0.6264 / N/A | 0.4458 / N/A | 0.2163 / N/A | 0.3730 / N/A |
| Cross-modal autoencoders | RMSE | 0.5725 / N/A | 0.3585 / N/A | 0.1917 / N/A | 0.2551 / N/A |
| BABEL | RMSE | 0.4335 / N/A | 0.3673 / N/A | 0.1816 / N/A | 0.2394 / N/A |
| scTAG | ARI | 0.75 / N/A | 0.96 / N/A | 0.53 / N/A | 0.60 / N/A |

**Table 8:** Modality Matching Task: performance comparison between our and original implementations in DANCE.

| Model | Evaluation Metric | GEX2ADT<br>(current/reported) | GEX2ATAC<br>(current/reported) |
| --- | --- | --- | --- |
| ScMoGCN | ACC | **0.0827** / 0.0810 | **0.0600** / 0.0630 |
| SCMM | ACC | 0.005 / N/A | 5e-5 / N/A |
| Cross-modal autoencoders | ACC | 0.0002 / N/A | 0.0002 / N/A |

**Table 9:** Joint Embedding Task: performance comparison between our and original implementations in DANCE.

| Model | Evaluation Metric | GEX2ADT<br>(current/reported) | GEX2ATAC<br>(current/reported) |
| --- | --- | --- | --- |
| ScMoGCN | ARI | 0.706 / N/A | 0.702 / N/A |
| ScMoGCNv2 | ARI | **0.734** / N/A | N/A / N/A |
| scMVAE | ARI | 0.499 / N/A | 0.577 / N/A |
| scDEC(JAE) | ARI | 0.705 / N/A | **0.735** / N/A |
| DCCA | ARI | 0.35 / N/A | 0.381 / N/A |

**Table 10:** Spatial Domain Task: performance comparison between our and original implementations in DANCE.

| Model | Evaluation Metric | Slice 151673<br>(current/reported) | Slice 151676<br>(current/reported) | Slice 151507<br>(current/reported) |
| --- | --- | --- | --- | --- |
| SpaGCN | ARI | 0.51 / 0.522 | 0.41 / N/A | 0.45 / N/A |
| STAGATE | ARI | **0.59** / N/A | **0.60** / 0.60 | **0.608** / N/A |
| stLearn | ARI | 0.30 / 0.36 | 0.29 / N/A | 0.31 / N/A |
| Louvain | ARI | 0.31 / 0.33 | 0.2528 / N/A | 0.28 / N/A |

**Table 11:** Cell Type Deconvolution: performance comparison between our and original implementations in DANCE.

| Model | Evaluation Metric | GSE174746<br>(current/reported) | CARD Synthetic<br>(current/reported) | SPOTlight Synthetic<br>(current/reported) |
| --- | --- | --- | --- | --- |
| DSTG | MSE | 0.172 / N/A | 0.0247 / N/A | 0.042 / N/A |
| SpatialDecon | MSE | 0.0014 / 0.009 | **0.0077** / N/A | **0.0055** / N/A |
| SPOTlight | MSE | 0.0098 / N/A | 0.0246 / 0.118 | 0.0109 / 0.16 |
| CARD | MSE | **0.0012** / N/A | 0.0078 / 0.0062 | 0.0076 / N/A |

## C Details of Supported Datasets

All supported datasets across 8 tasks in DANCE are summarized in Table 12. For each supported dataset, we list what type of species and tissue it is about, dataset dimensions including the number of cells and genes, and also the protocol about how to generate the dataset for reference. In the column of “Availability”, the dataset link is provided once you click the reference.

**Table 12:**
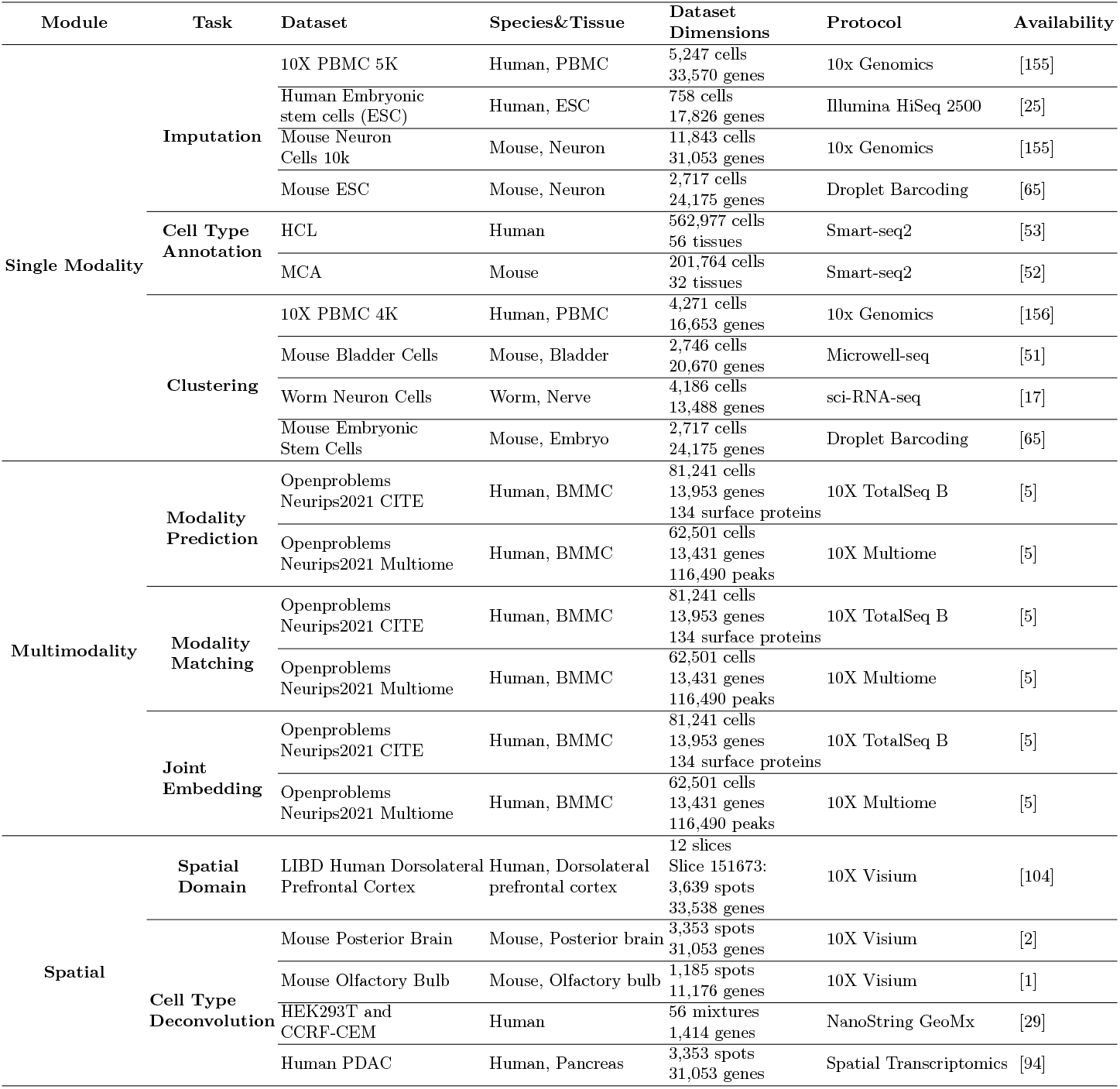
A summary of all supported datasets in DANCE.

## D Details of Supported Tasks and Models

## D.1 Single Modality Module

## Imputation

The goal of imputation for scRNA-seq data is to address artificial zeros in scRNA-seq data generated during the sequencing process systematically or by chance due to technological limitations. Imputation aims at correcting these artificial zeros by filling in realistic values that reflect true biological gene expressions [71]. Thus, a good imputation method should be able to distinguish artificial zeros from biologically true zeros and recover true expressions for artificial zeros. As the corresponding biologically true expression values are unavailable for entries of artificial zeros in the gene-cell matrix, dropouts are simulated for benchmarking such that metrics such as cosine similarity, correlations, or MSE-related metrics can then be used to evaluate imputation algorithms; alternatively, prior cell information, such as cell types, can be used to evaluate whether an imputation method could recover biological signals by comparing clustering patterns using original and imputed expressions [6, 31, 56, 75, 114, 139].

## dance.modules.single_modality.imputation.deepimpute

DeepImpute [6] builds multiple neural networks in parallel to impute target genes using a set of input genes. Given a scRNA-seq matrix *X*, target genes, i.e, genes to be imputed, are selected based on the variance over mean ratio, which are split into *N* random subsets. Each subset corresponds to a neural network consisting of two layers: a dense layer and a dropout layer with ReLu and softplus as activations, respectively. The inputs to each neural network are genes highly correlated with corresponding target genes based on Pearson’s correlation coefficient. The loss function is the Weighted mean squared error (MSE).

## dance.modules.single_modality.imputation.scgnn

scGNN [139] uses an integrative autoencoder framework for scRNA-seq gene expression imputation that incorporates gene regulatory signals (TRS). It includes a feature autoencoder, a graph autoencoder, and a cluster autoencoder that are trained iteratively, whose outputs are used in the final imputation autoencoder to recover gene expressions.

scgnn uses left-truncated mixed Gaussian (LTMG) to account for regulatory signals. The normalized expression for a gene is modeled by a mixture of *k* Gaussian distributions representing *k* TRSs, and left truncation is used to account for dropouts and lowly expressed values. The expression of gene *i* in cell *j* can then be assigned to the TRS under whose Gaussian distribution the observed value is most likely to occur. All parameters in this step are estimated by MLE.

Given an input scRNA-seq expression matrix *X*, the feature autoencoder extracts a lower-dimensional embedding *X′* and reconstructs the expression 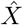, which is composed of two dense layers in both the encoder and the decoder. The loss function is MSE integrated with gene regulation information

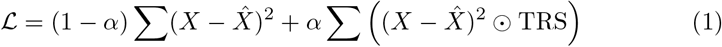

where *α* ∈ [0, 1] and ⊙ denotes element-wise product.

A KNN cell graph is constructed from the learned embeddings in the feature autoencoder, which is pruned by the Isolation Forest. With node feature matrix *X′*, the two-layer GCN encoder is constructed as

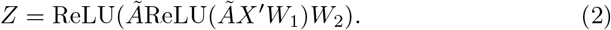

where *Ã* is the symmetrically normalized adjacency matrix of the cell graph and *W*_1_, *W*_2_ are learnable parameters.

The associated decoder is:

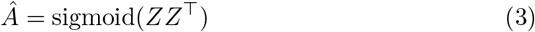

The parameters are learned through the cross-entropy loss

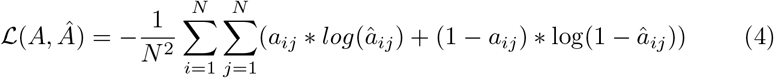

where *a_ij_* and *â_ij_* are elements in A and *Â*.

The *k*-means clustering is then applied to the learned embedding in the graph autoencoder. The number of clusters is determined by the Louvain algorithm. Each cell cluster has an autoencoder to regenerate gene expressions within a cluster, whose structure is the same as the feature autoencoder without being regularized by TRS. The reconstructed gene expression is fed back into the feature encoder iteratively until the convergence of adjacency matrix (*Ã_t_* − *Ã*_*t*−1_ < *γ*_1_)and cell clusters converge (ARI> *γ*_2_).

After convergence, a final imputation autoencoder is trained, which has a similar structure and loss to the feature autoencoder but with three additional regularizations:

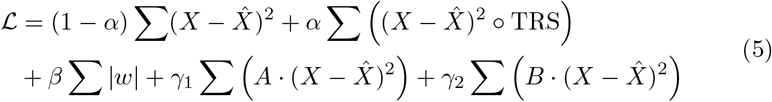

where *β* ∈ [0, 1] controls the intensity of the *L*1 penalization and *B* is a matrix indicating whether two cells belong to the same cluster.

## dance.modules.single_modality.imputation.graphsci

GraphSCI [114] is a GNN-based method to impute scRNA-seq data expressions. Given a gene expression matrix *X* with *N* genes and *M* cells, a gene graph associated with an *N* × *N* adjacency matrix *A* is derived based on gene-wise Pearson correlation coefficients. The gene expression matrix and the constructed gene graph are the input to 1) a two-layer GCN whose lower-dimensional embedding gives the Gaussian distribution representing gene relationships and 2) a two-layer fully connected neural network (NN) from which a ZINB distribution for scRNA-seq data expressions can be inferred. In particular, the GCN is formulated as

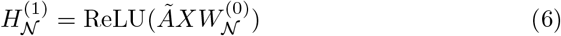

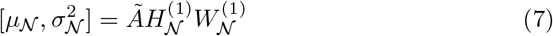

where 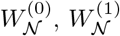 are the trainable parameters.

The fully-connected neural network is defined as

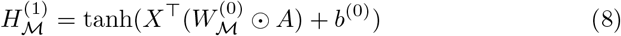

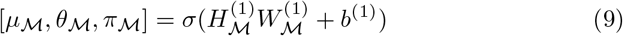

where 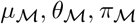 are the mean, dispersion, and dropout probability for the ZINB distribution, and 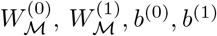 are trainable parameters.

The learned latent variables 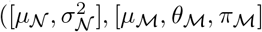 are re-parameterized as 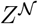 and 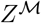, i.e, the latent space representation of the GCN and NN, to be used by the decoder to construct the imputed gene expression 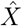. Specifically, the imputed gene expression for gene *i* in cell *j* is

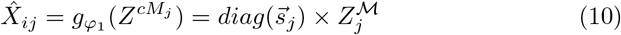

where 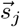 is the size factor of cell *j*.

Meanwhile, the reconstructed edge weight between gene *i* and gene *j* is

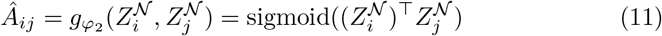

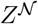 and 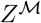 are optimized by variational lower bound

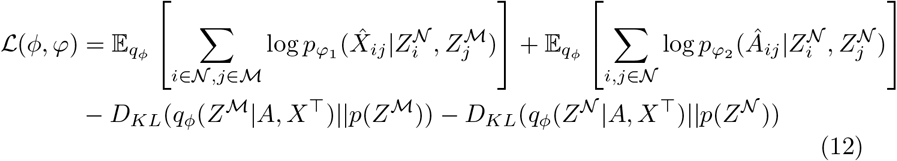

where 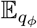 is the cross-entropy function and and *D_KL_*(*q*| | *p*) is the Kullback-Leibler divergence between distributions *q* and *p*.

## Cell Type Annotation

Cell type annotation targets applying statistics of cellular properties to infer cell types. Given the gene expression of several cell types, for each cell with a certain single-cell expression matrix, the degree of similarity can be calculated. Based on the optimal similarity result, the cell type can then be inferred. In DANCE, we support 5 models that establish measurements of evaluating the similarity of gene expression profiles of unknown cells to gene expression matrices of known cell types. The model performance is evaluated by prediction accuracy.

## dance.modules.single_modality.cell_type_annotation.scdeepsort

Scdeep-sort [118] includes three modules: an embedding layer, a weighted graph aggregator, and linear classification towers. Given *m* genes and *n* cells, we have the input single-cell data matrix *D* ∈ ℝ^*m*×*n*^. To generate the weighted cell-gene graph, we first apply PCA to extract *d* dimensional representations of initial representations. A weighted adjacency matrix 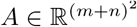 and a node embedding *X* ∈ ℝ^(*m*+*n*)×*d*^ are generated from *D*. For a gene node *j*, the shareable parameter *β_j_* denotes the confidence value for the edges interacting with node *j*. For the self-loop edge for each cell, we use *α* to denote its confidence value. Let 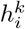 be the node *i*’s embedding vector in the *k_th_* layer, the aggregator layer is

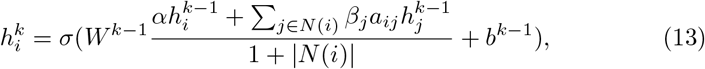

where *a_ij_*is the normalized weight of an edge from nodes *i* to *j*. Then cell node representations are fed into linear classifier layers

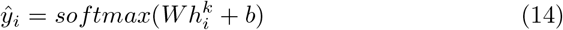

Cross entropy loss is applied in the objective function:

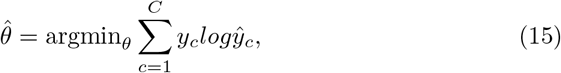

where *θ* represents all trainable parameters.

## dance.modules.single_modality.cell_type_annotation.celltypist Celltypist

[32] uses a multinomial logistic regression classifier defined as:

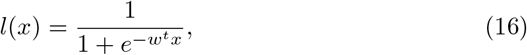

where *l*(*x*) is the decision score with *x* as the input vector. Then the decision score for each cell is defined as the linear combination of the scaled gene expression and the model coefficients associated with the given gene type, and the possibility is calculated by transforming the decision score by a sigmoid function.

## dance.modules.single_modality.cell_type_annotation.singlecellnet

Single-Cellnet [128] revamped the random forest classifier method to enable classification of scRNA-seq data cross platforms and cross-species. It sends the input features into a number of decision tree classifiers and uses majority voting to make predictions.

## dance.modules.single_modality.cell_type_annotation.actinn

ACTINN [81] proposes a neural network based model for cell type annotation. It applies multilayer perceptron for the identification of cell types that can be implemented as:

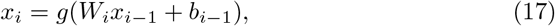

where *x_i_* is the output from the *i_th_* layer, *W_i_* and *b_i_* represent the weight matrix and the bias in the *i_t_h* layer and *g* represents the activation function used in the neural network. In the input layers and hidden layers, the activation function is ReLU as:

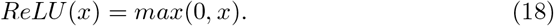

It utilizes the softmax function as the activation function *g* for the output layer, which is defined as:

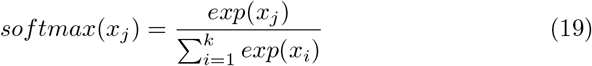

where *x_j_* represents the *j_th_* element of the input vector for the output layer and *k* is the length of the vector *x*.

## dance.modules.single_modality.cell_type_annotation.svm

Support vector machine(SVM) is widely adopted as a benchmark in many studies [3,118]. Given the input *x* and label *y*, the prediction function is

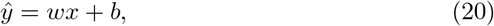

where *w* represents the weights and *b* is the interception in the SVM. Based on Eq 20, SVM optimizes the following problem:

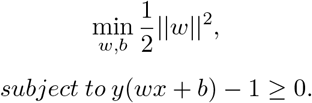

## Clustering

Clustering is a crucial part of single-cell analysis. With clustering, researchers can identify cell types or cell-type subgroups within the gene expression data. In the clustering task, we now support 5 models. The first 3 models are GNN based, and the later 2 models are non-GNN based with autoencoder as the backbone. The clustering performance is evaluated by Adjusted Rand index (ARI).

## dance.modules.single_modality.clustering.scdeepcluster

scDeepCluster [131] introduces a ZINB-based autoencoder. The input matrix is corrupted by a Gaussian noise *e*: *X*^corrupt^ = *X* + *e*. Then the encoder produces latent representation *Z* from *X*^corrupt^. The decoder can be formulated as:

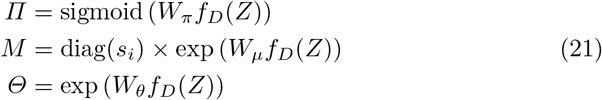

where *f_D_*(·) is the decoder function; *s_i_* is the size factor; {*Π*, *M*, *Θ*} are the estimations of ZINB distribution parameters {*π*, *μ*, *θ*} respectively. The clustering process is based on soft assignment. The soft label *q_ij_* of embedded point *z_i_* is defined as:

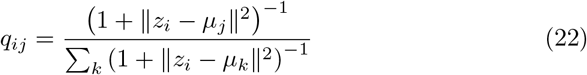

where *μ_j_* is the *j*-th cluster center.

## dance.modules.single_modality.clustering.scdcc

scDCC [132] shares the same model structure as scDeepCluster. In the training process, pairwise constraints are integrated into the loss function. There are two types of pairwise constraints, i.e., must-link (ML) and cannot-link (CL). Two instances with a must-link constraint should have similar soft labels as:

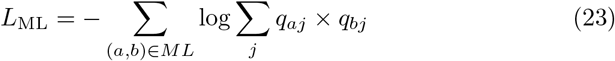

While two instances with cannot-link should have different soft labels as:

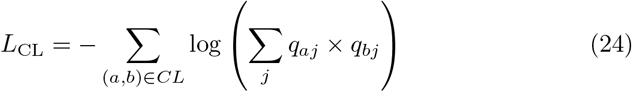

where *q_aj_* and *q_bj_* are soft labels defined in (22).

## dance.modules.single_modality.clustering.graphsc

graph-sc [26] utilizes gene-to-cell graph as the input of graph autoencoder. In the gene-to-cell graph, genes and cells are nodes, and there are weighted edges between cell nodes and the expressed gene nodes. Let the raw data matrix be *X*, then the weight of gene *i* to cell *j* is 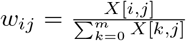. We use *W*, *Z*_0_, and *A* to denote the graph weight matrix, the input features, and the adjacency matrix. 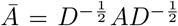 is the normalized adjacency matrix with *D* as the degree matrix of *A*. The cell embeddings can be obtained by:

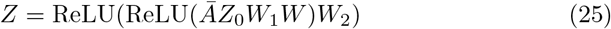

where *W*_1_ and *W*_2_ are learnable weights. The new adjacency matrix *Â* is then reconstructed by *Â* = sigmoid(*ZZ*^⊤^).

## dance.modules.single_modality.clustering.sctag

scTAG [151] first generates a K-nearest neighbor cell-to-cell graph. It then adopts a ZINB-based graph autoencoder to process it, which takes topology adaptive graph convolutional network (TAGCN) [35] as the graph encoder. Consider the *l*-th hidden layer, let the input data be 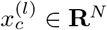 where *c* = 1, 2, *C_l_*, *C_l_* is the number of features of each node, and *N* denotes the number of samples. The graph convolution process can be defined as follows:

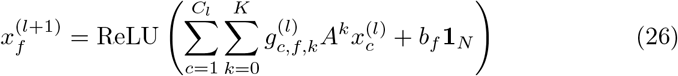

where *b_f_* is a learnable bias; *K* is the number of convolution kernels; and 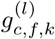 denotes the polynomial coefficients. Denote *Z* as the latent embedded representation. The new adjacency matrix *Â* is then reconstructed by *Â* = sigmoid(*ZZ*^⊤^). The estimations of ZINB distribution parameters {*π*, *μ*, *θ*} are obtained by (21).

## dance.modules.single_modality.clustering.scdsc

scDSC [42] consists of a ZINB-based autoencoder and a graph autoencoder with the KNN cell-to-cell graph. In the ZINB-based autoencoder, let the latent representation be *H*, the output of last decoding layer be *D*, *b*_enc_ and *b*_dec_ be bias of encoder and decoder, respectively. The autoencoder is formulated as follows:

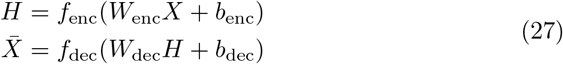

where 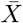 is the reconstructed expression matrix. The estimations of ZINB distribution parameters {*π*, *μ*, *θ*} are similar to those in (21). In the graph encoder, denote the representation of *l*-th layer as *Z_(l)_*, the adjacency matrix as *A*, and the degree matrix as *D*. The new representation is generated by:

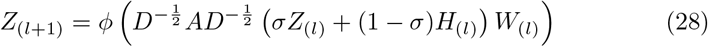

where *ϕ*(·) is an activation function; *σ* is a hyperparameter; and *H_(l)_* is the representation of *l*-th layer of ZINB-based encoder.

## D.2 Multimodality Module

## Modality Prediction

Modality prediction is to predict features of a target modality from features of an input modality. The evaluation is based on rooted mean squared error (RMSE) between ground-truth features and prediction. In this task, DANCE supports 4 models. All of them are deep learning models, one of which is based on graph neural networks.

## dance.modules.multi_modality.predict_modality.scmogcn

scMoGNN [143] first converts the input feature matrix into a cell-feature bipartite graph 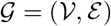 where each node *v* represents a cell or a feature. For instance, for the input of gene expression, each node can be a cell or a gene. Meanwhile, additional gene-gene connections are added based on an external pathway dataset [127]. Every pair of vertices in *V* are connected by a weighted edge, which either depends on the read count of a feature in a cell, or the correlation between features.

Specifically, we use **X** ∈ ℝ^*N*×*k*^ to denote the feature matrix of input modality with *N* the number of cells and *k* the dimension of input features. The adjacency matrix **A** of the constructed cell-feature bipartite graph 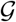 is:

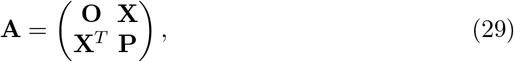

where **O** is a zero matrix, and **P** ∈ ℝ^*k*×*k*^ indicates the gene-gene links.

To learn node embeddings on 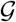, scMoGNN introduce a heterogeneous graph convolutional network. The proposed heterogeneous graph convolutional network can be stated as:

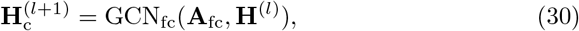

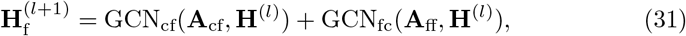

where 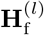 is the embeddings of feature nodes in *l*-th layer, 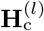 is the embeddings of cell nodes in *l*-th layer. Subscripts cf, *textfc* and *textff* denote the graph convolution over all the cell-to-feature edges, feature-to-cell edges and feature-to-feature edges respectively. After stacking convolutional layers, cell node embeddings from each layer are collected by a weighted sum, denoted as

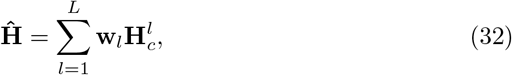

where **w** is a learnable weight vector, *L* is the total number of layers. **Ĥ** is then passed to the downstream modules. In the case of modality prediction, a fully-connected predictive head is added. The final prediction **Z** ∈ ℝ^*N*×*d*^ of scMoGNN can be thus written as:

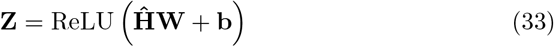

where **W** and **b** are the parameters of the predictive head. Overall, a mean squared error MSE(**Z**, **Y**) is optimized through training, where **Y** ∈ ℝ^*N*×*d*^ is ground-truth features of the target modality.

## dance.modules.multi_modality.predict_modality.babel

BABEL [146] trains two neural-network-based encoders and two decoders on the paired data to translate data from one modality to the other and to reconstruct itself, thus eventually obtaining shared embedding. A special design in BABEL is to add a prior distribution to the decoder. Instead of directly outputting the feature values, the decoder estimates the parameters of feature distribution.

Formally, for each cell, the RNA decoder models the likelihood of expressions *y* as a negative binomial (NB) distribution, written as:

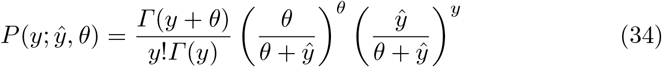

where *Γ* denotes the gamma function, *ŷ* and *θ* are the estimation of the mean and dispersion of the distribution. In practice, these estimations come from neural network encoders and decoders. The optimization problem is thus formalized by minimizing the negative log-likelihood:

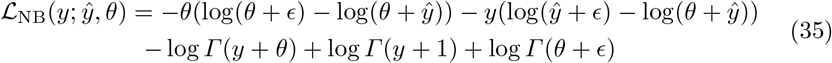

where *ϵ* is a tiny constant for numerical stability.

The ATAC decoder has a different modeling approach since the feature for each peak is binary. The loss function for the ATAC decoder is based on binary cross-entropy, shown below:

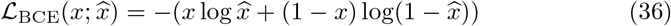

where *x* represents ground-truth ATAC-seq features, and 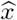 denotes the prediction from the ATAC decoder.

The overall loss function is hereby formulated as:

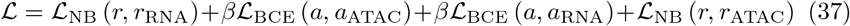

where *r* and *a* are the ground-truth RNA-seq features and ATAC-seq features respectively, subscripts indicate from which modality the features are predicted (e.g., *a*_RNA_ represents the ATAC features predicted from RNA). The first two terms in the loss function are similar to reconstruction loss in autoencoders. The latter two terms can be considered as modality prediction loss.

For testing, we simply take *a*_RNA_ as the prediction of ATAC-seq from RNA-seq, and take *r*_ATAC_ as the prediction of RNA-seq from ATAC-seq.

## dance.modules.multi_modality.predict_modality.cmae

Cross-modal Autoencoders [149] uses autoencoders to map vastly different modalities (including images) to a shared latent space. Specifically, a discriminator and adversarial loss are added to force the distributions of different modalities to be matched in the latent space. To make use of prior knowledge, an additional loss term can further be added to align specific markers or anchoring cells.

Formally, an invariant latent distribution for two modalities *i* and *j* is learned as follows. We denote the input features of two modalities as **X**_*i*_ and **X**_*j*_ For modality *i*, we optimize the objective:

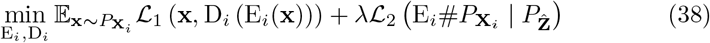

while for modality *j*, we optimize:

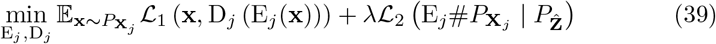

Here, 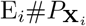 refers to the distribution of modality *i* in the latent space *Z*, 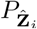 is the expected distribution of joint latent space *Z*. D and E refer to encoders and decoders parameterized by neural networks. 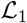 is the Euclidean distance metric, which is equivalent to a reconstruction loss. 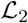 represents a divergence between probability distributions, since 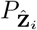 is unknown, it can be adapted to a discriminator and an adversarial loss.

Several additional losses can be added to the model to incorporate prior knowledge. For example, if training data include paired multimodal data, those cells with more than one modality can be considered as anchors. Suppose 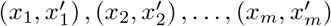 are corresponding points from two datasets, we can add the following anchor loss,

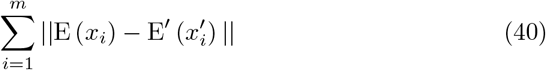

where E and E′ are encoders for the two modalities, respectively.

The final prediction from modality *i* to modality *j* is *D_j_* (E_*i*_(**X**)).

## dance.modules.multi_modality.predict_modality.scmm

scMM [92] leverages a mixture-of-experts (MoE) multimodal variational autoencoder [119] (VAE) to explore the latent dimensions that associate with multimodal regulatory programs. It models raw count features from each modality using various probability distributions in an end-to-end way. Specifically, an MoE multimodal VAE (MM-VAE) is to learn a multimodal generative model:

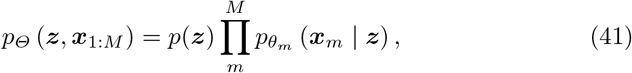

where 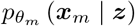 is the likelihood for *m*-th modality, and 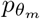 is parameterized by a neural network decoder. To optimize the model, a typical training objective for VAE is to maximize the ELBO:

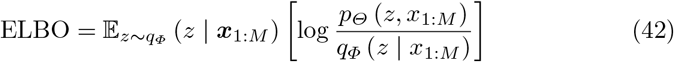

where *q_Φ_* (***z*** | ***x***_1:*M*_) is the joint variational posterior that can be parameterized by a neural network encoder. In addition, an MMVAE factorizes the joint posterior with an MoE:

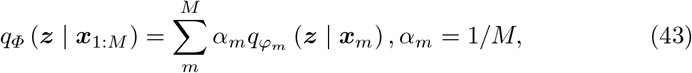

where the posterior of the *m*-th modality is 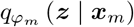. It is parameterized by the encoder. When using stratified sampling, ELBO can be re-written as:

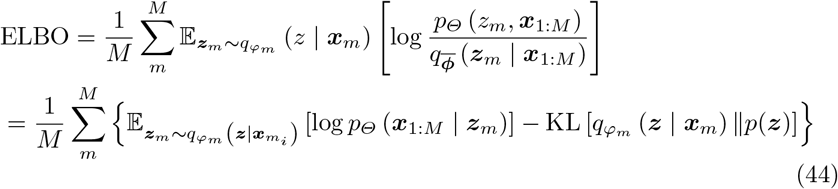

The first expectation term is to measure the reconstruction performance. Here, latent variables of each modality would be used to reconstruct all modalities, including cross-modal translation. The second term is a regularization term forcing the variational posterior to be consistent the prior distribution *p*(***z***), which is a Laplacian distribution in practice. In the modality prediction task, we take the cross-modal generation results as the prediction of the model.

## Modality Matching

The objective of the modality matching task is to identify the cell correspondence across modalities. To be concrete, we separate each modality of the jointly profiled dataset into a subset, and the order of cells in each subset is disturbed. In the training dataset, the cell correspondence labels between subsets are given. While in the testing data, the correspondence is not given. The model needs to learn to identify cell correspondence from the labeled training data and evaluate it on the testing data.

To provide a more flexible protocol, the model output is adapted to a matching score matrix **S** ∈ ℝ^*n*×*n*^, where *n* is the number of cells, **S**_*i,j*_ is the probability that cell *i* from one modality corresponds to cell *j* from the other modality. Therefore, **S** is a non-negative matrix where each row sums to 1. As metrics, we compute the average probability assigned to the correct matching. In this module, DANCE now supports 3 models. All of them are deep learning models, one of which is based on graph neural networks.

## dance.modules.multi_modality.match_modality.scmogcn

The overall structure of scMoGNN in the modality matching task is the same as in the modality prediction task. However, in the modality prediction task, the input is only one modality, while in the modality matching task, features of two modalities are given altogether. Therefore, scMoGMM constructs two graphs 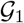 and 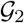 for two modalities respectively. The cell node embeddings are obtained in the same way as before, denoted as **Ĥ**_1_ and **Ĥ**_2_. Then a matching head is added, formulated as:

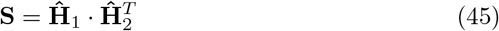

where **S** is the desired output matrix of matching scores. For training, we calculate the cross entropy loss between **S** and a ground-truth matching matrix **M** ∈ ℝ^*n*×*n*^, where **M**_*i,j*_ = 1 only when cell *i* in modality 1 corresponds to cell *j* in modality 2. In addition, several auxiliary losses are added, including a reconstruction loss and a translation loss.

For testing, a bipartite matching via the Hungarian algorithm is implemented as post-processing. It generates an optimized sparse score matrix over the raw **S** matrix.

## dance.modules.multi_modality.match_modality.cmae

The overall structure of Cross-modal Autoencoders is the same as in the modality prediction task, where we implement encoders and decoders for all the modalities. Hereby in the modality matching task, we directly utilize the latent space instead of using a decoder to generate target modality.

To be specific, the embeddings from each modality can be denoted as:

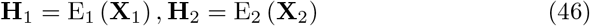

where we denote input matrix, encoder and embeddings of *m*-th modality as E_*m*_, **H**_*m*_ and **X**_*m*_ respectively. Then the score matrix is obtained by:

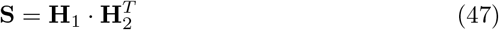

where **S** is the output score matrix.

## dance.modules.multi_modality.match_modality.scmm

The overall structure of scMM is the same as in the modality prediction task, where we implemented a neural network encoder E_*m*_ for each modality *m* to estimate the variational posterior *q_Φ_* (***z*** | ***x***_1:*M*_). In the modality matching task, we hereby take the latent vectors generated by encoders as the source for matching. The whole process is the same as Eq. 46 and 47.

## Joint Embedding

Joint embedding aims to encode features from two modalities into a low-dimensional joint latent space. To be consistent with the NeurIPS competition [80], we set the latent dimension size to 100. For the evaluation, currently, we only support NMI and ARI with the k-means clustering as metrics in our DANCE package. These metrics evaluate the consistency between latent clusters and the ground-truth cell type labels. More comprehensive metrics were introduced in the competition, and we are going to incorporate them into our package in the future. In this module, we now support 4 models. All of them are deep learning models, one of which is a graph neural network.

## dance.modules.multi_modality.joint_embedding.scmogcn

The overall structure of scMoGNN in the joint embedding task is still similar to what is shown in the modality prediction task. However, different from previous tasks, here scMoGNN first preprocesses data as suggested by Seurat [125]. It reduces the dimension of modality *m* to a predefined *k_m_* dimension, using latent semantic indexing (LSI). Empirically, we set *k_m_* = 256 for RNA-seq features, *k_m_* = 512 for ATAC-seq features, and no dimension reduction for surface protein features. Next, the preprocessed features of two modalities are concatenated and jointly considered as feature nodes in the graph construction, as described in Eq. 29. Since no feature interactions are specified, here we replace matrix **P** in Eq. 29 with **O**. With the graph 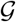 so constructed, scMoGNN is further trained by minimizing a reconstruction loss, a cell type auxiliary loss and a regularization loss.

Specifically, cell embeddings **Ĥ** are obtained as Eq. 32. An MLP decoder *f_θ_* is involved to reconstruct the input features from **Ĥ**. The first *T* dimensions in **Ĥ** are also used to predict cell type, where *T* is equal to the number of predefined cell types, and a regularization loss is added to the rest of the dimensions. The overall loss function can be written as:

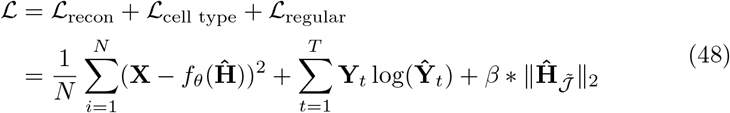

where *f_θ_* is a two-layer MLP decoder, 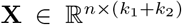 is the pre-processed feature matrix, *k*_1_ and *k*_2_ are specified feature dimensions of two modalities, **Y** ∈ ℝ^*N*×*T*^ is predefined cell types for each cell in a sparse form, and 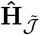 refers to the hidden dimensions other than first *T* dimensions. **Ŷ** is calculated by a softmax function over the first *T* dimensions of **Ĥ** formulated as:

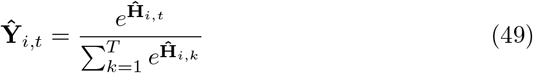

In the end, **Ĥ** is the resulting joint cell embeddings from scMoGNN, which is expected to encode cellular information that is essential in the joint embedding task.

## dance.modules.multi_modality.joint_embedding.jae

JAE is an adapted model from scDEC [78]. It is proposed by the authors of scDEC in the NeurIPS competition [80] to better leverage cell annotations. Formally, JAE follows the typical autoencoder architecture with an encoder *f_θ_* and a decoder *g_θ_*. They are parameterized by MLPs, denoted as:

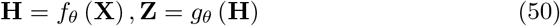

where **H** ∈ ℝ^*n*×*d*^ is the joint embeddings, and **Z** is the recovered input features. The novelty of JAE is that it separates cell embeddings into four parts, written as:

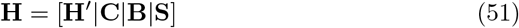

where | denotes concatenation, **C** ∈ ℝ^*n*×*c*^ is additionally supervised by cell type labels, **B** ∈ ℝ^*n*×*b*^ is additionally supervised by batch labels, **S** ∈ ℝ^*n*×*s*^ is additionally supervised by cell cycle phase score, **H′** ∈ ℝ^*n*×*z*^ is the remaining dimensions. Therefore *z* + *b* + *s* + *c* = *d*. The overall loss function of JAE is formulated as:

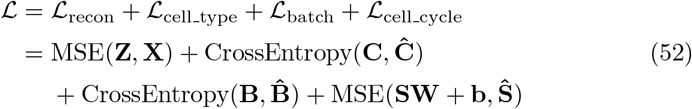

where **Ĉ**, 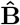, **Ŝ** are cell type labels, batch labels and cell cycle scores, respectively. **W** ∈ ℝ^*s*×2^ and **b** ∈ ℝ^2^ are an extra linear transformation to project **S** to cell cycle score vector space. Eventually, **H** is the output joint embedding from JAE.

## dance.modules.multi_modality.joint_embedding.scmvae

scMVAE [157] learn the distribution of multi-omics via three learning strategies simultaneously: product of experts (PoE), neural networks, and concatenation of multi-omics features. In addition, scMVAE models raw count features from each modality through a ZINB distribution. Specifically, let *z* be the joint embeddings obtained from multimodal encoders. *p*(*z*|*c*) is a Gaussian mixture distribution. Its mean vector *μ_c_* and a covariance matrix *σ_c_* are conditioned on *c*, where *c* is a discrete categorical variable indicating cell types. This Gaussian mixture prior of *z* is introduced in MVAE to enhance the interpretability of the latent representations and the performance of generation. Let *x* and *y* be features of two modalities. Then the distribution *p*(*x*, *y*, *z*, *c*, *l_x_*, *l_y_*) is formulated as:

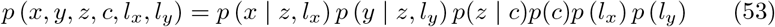

where *l_x_* and *l_y_* are the library size factors of two modalities.

The prior distributions for all the random variables are listed below:

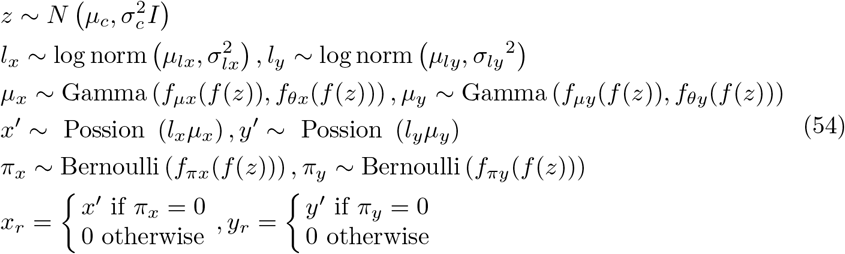

where *f_θx_* (*f*(*z*)) and *f_θy_* (*f*(*z*)) are the inverse desperations of two modalities from the variational Bayesian inference. *f_μx_* and *f_μy_* are two neural network decoders that estimate the mean proportions of features for two modalities in each cell by using a softmax function, which simulates the library-size normalized features. *f_πx_* and *f_πy_* are neural network decoders that estimate the probability of features being dropped out due to technical issues. They use a sigmoid function to model this probability.

To train scMVAE, we maximize the log-likelihood of the multi-omics observations. Following the convention of variational autoencoders, this objective is converted to optimizing an evidence lower bound (ELBO):

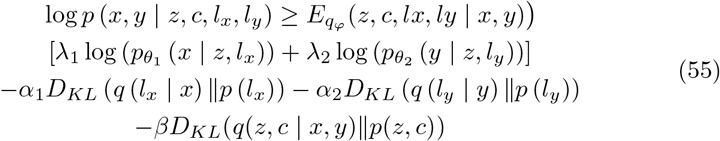

Both modalities in the ELBO have two reconstruction terms, and three regularization terms are implemented by KL divergence. *q_φ_* refers to a multimodal encoder. While 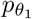 and 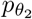 refer to decoders, for two modalities respectively. The latent vector *z* estimated from E is the eventual joint embedding.

## dance.modules.multi_modality.joint_embedding.dcca

In DCCA [157], data of each modality are modeled by a variational autoencoder (VAE). Specifically, for modality *m*, an encoder *E_m_* transforms the input features into latent space *z_m_*. A decoder *D_m_* then transforms *z_m_* into the parameters of the NB or Bernoulli. For example, RNA-seq and ADT data follow NB distribution, denoted as:

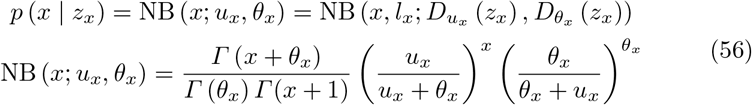

where 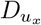 and 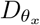 are decoders, each dimension of *u_x_* and *θ_x_* indicates the mean and variance of NB distribution for each feature, and one-dimensional constant variable *l_x_* indicates the library size of each cell. While ATAC-seq data follow Bernoulli distribution, denoted as:

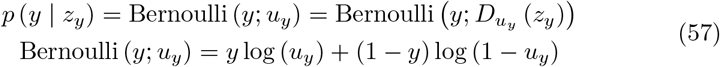

where *D_uy_* refers to the ATAC-seq decoder.

Each VAE is first trained separately with each modality. Then, two VAEs are trained together to maximize the similarity between two latent spaces. For example, given the embeddings from a VAE_RNA_ and a VAE_ATAC_, they optimize an objective function that combines reconstruction loss with the cell embeddings similarity loss. Hence, the total ELBO for VAE_RNA_ and VAE_ATAC_ can be written as:

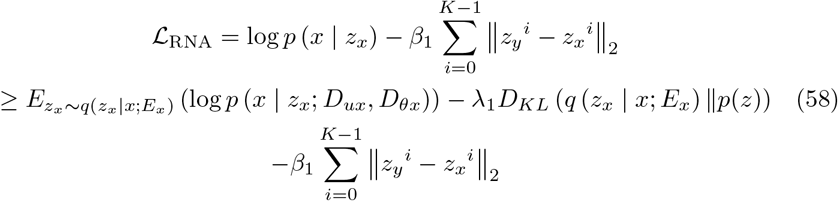

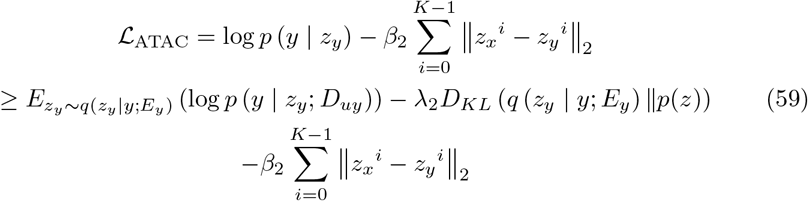

According to the DCCA paper, after jointly training two VAEs, the embedding from VAE_RNA_ is selected as the final embedding for downstream analysis.

## D.3 Spatial Transcriptomics Module

## Spatial Domain

In spatial transcriptomics, the spatial data is referring to spots with x,y coordinates, and each spot captures several cells. The objective of the spatial domain is to partition the spatial data into meaningful clusters. Each cluster uncovered by this analysis is regarded as a spatial domain. Spots in the same spatial area are comparable and consistent in gene expression and histology, but spots in different spatial regions are distinct [13]. For evaluation, Adjusted Rand Index (ARI) [150] is utilized to compare the efficacy of various clustering techniques. It computes the similarity between the algorithm-predicted clustering labels and the actual labels. In the spatial domain task, DANCE supports 4 models including 2 GNN-based models and 2 traditional models.

## dance.modules.spatial.spatial_domain.spagcn

SpaGCN [57] first constructs a weighted undirected graph, *G*(*V*, *E*) from the gene expression and histological image data. In *G*, each vertex *v* ∈ *V* is a spot, and every pair of vertices in *V* are connected by a weighted edge, which assesses the correlation between the two spots. The weight for the pair (*u*,*v*) is calculated as:

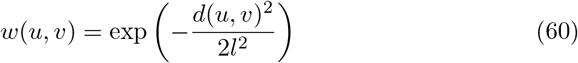

where hyperparameter *l* represents characteristic length scale, and *d*(*u*,*v*) calculates Euclidean distance between spots *u* and *v*. This distance is computed by the spatial distance and the corresponding histology information between two spots. The initial node representation in the graph is gene expression after dimension reduction. A process known as graph convolution is utilized by SpaGCN to aggregate gene expression data in accordance with edge weights. Then the output of the graph convolution layer would be new node representation capturing information on gene expression, histology and physical location. Based on the new spot representation, an unsupervised clustering algorithm is further employed to iteratively cluster the spots into spatial domains. The probability of assigning spot *i* to cluster *j* is defined as:

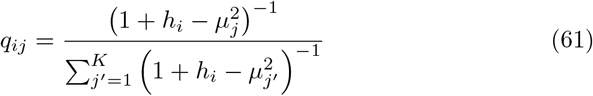

where *h_i_* is the embedded point for spot *i*, and *u_j_* indicates centroid *j*. Then the clusters are refined iteratively by a target distribution *P* from *q_ij_*:

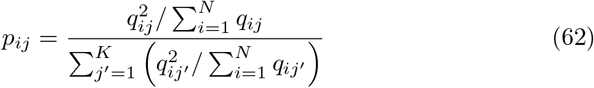

which gives more weight to locations that have been recognized with high confidence and normalizes each centroid’s contribution to the loss function to avoid having large clusters distort the hidden feature space. The objective function is based on a Kullback–Leibler (KL) divergence as:

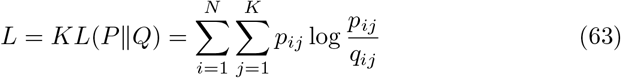

where *N* is the number of spot samples, and *K* is the number of clusterings.

## dance.modules.spatial.spatial_domain.stagate

STAGATE [33] is a graph attention based autoencoder [116] with encoder, decoder and graph attention layers. For graph construction, it builds up an undirected graph with a radius *r* that has been predefined based on physical distance. We use**A** to denote the adjacency matrix of the graph where **A_ij_** = 1 if the Euclidean distance between spots *i* and *j* is less than *r*. In addition, STAGATE also builds up a cell type-aware graph via updating the previously constructed graph based on the pre-clustering of gene expressions.

The encoder in STAGATE takes the gene expressions that have been normalized as its inputs. It then generates embedding for each spot by aggregating information from its surrounding nodes collectively. The latent representation for spot *i* is calculated as:

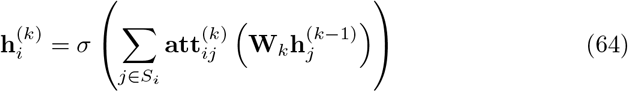

where *σ* is the activation function, **W**_*k*_ is the trainable weight matrix, *S_i_* is the neighbors of spot *i*, *k* indicates *k*-th encoder layer and 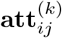 is the attention score (i.e., the edge weight) between spots *i* and *j* obtained from the *k*-th graph attention layer’s output.

Contrarily, using the encoder’s output as input, the decoder transforms the latent embedding into a reconstructed normalized expression profile.

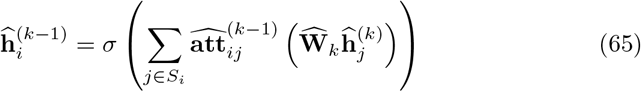

In graph attention layer, a widely-used self-attention technique for graph neural networks is adopted to learn the similarity between surrounding spots in an adaptive manner. The edge weight between spot *i* and its surrounding spot *j* in the *k*-th encoder layer is calculated as:

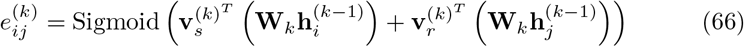

where 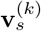 and 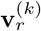 are the trainable weights.

The attention score is further normalized by a softmax function in the following way:

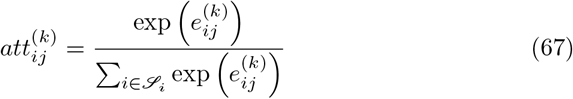

Eventually, STAGATE aims to minimize the reconstruction loss of normalized expressions in the following way:

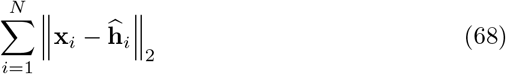

where **x**_*i*_ is the original normalized gene expressions, and 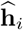 is the reconstructed normalized gene expression for spot *i*.

## dance.modules.spatial.spatial_domain.louvain

Louvain [11] is to extract the community structure of large networks, which draws inspiration from the optimization of modularity. A modularity measure shows how densely edges inside communities are clustered in comparison to edges outside communities. Theoretically, optimizing this value leads to the optimal grouping of nodes in a particular network. Due to the impracticality of traversing all possible iterations of the nodes into groups, heuristic approaches are utilized. The modularity is defined as:

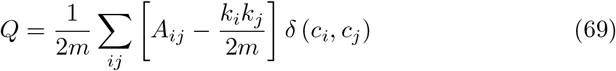

where *A_ij_* is the edge weight between nodes *i* and *j*, *m* represents the total weight of all edges in the graph, the weights of the edges that connect nodes *i* and *j* are added up to get *k_i_* and *k_j_*, *c_i_* and *c_j_* denote node communities, and *δ* represents Kronecker delta function [9] (*δ*(*x*, *y*) = 1 if *x* = *y*; else, 0).

To efficiently maximize the modularity, there are two looping phases. First, a community is associated with each node in the network. Because of this primary partitioning, there is consequently the same number of communities as nodes. Then, the change in modularity is determined for each node *i* by removing it from its own community and inserting it in the community of each of *i*’s neighbors *j*. Integration of a previously isolated node *i* into a community *C* results in an increase in modularity *ΔQ* equal to:

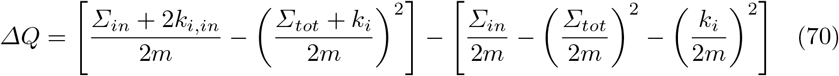

where ∑_*tot*_ is the total link weights occurring to nodes in *C*, ∑_*in*_ represents the total link weights within *C*, *k_i_* denotes the sum of the link weights associated with node *i*, *k_i,in_* represents the total link weights from node *i* to all other nodes in *C*, and *m* is the total link weights for the whole network. Once this value is computed for all communities to which *i* belongs, *i* is assigned to the community where the modularity increase was largest. If there is no possibility of expansion, *i* stays in the same community. This technique is repeated and applied successively to all nodes until no further growth in modularity is possible. The first step concludes when this modularity maximum is reached.

In the second phase, all nodes inside the same community are grouped together and a new network consisting of communities from the first phase is constructed. The connections between multiple nodes within the same community and a node in a separate community are now described by weighted edges between communities. The second stage is complete when the new network is set up, at which point the first stage can be applied to it again.

## dance.modules.spatial.spatial_domain.stlearn

stLearn [109] performs unsupervised clustering on SME-normalized data to group similar areas into clusters and discover sub-clustering alternatives based on the geographic separation of clusters inside the tissue. The name of this stLearn function is SMEcluster. Using normalized expression values, stLearn separates cell types in a tissue through a two-step spatial clustering approach. stLearn implements a conventional Louvain clustering technique for scRNAseq data as the initial step. Linear Principal Component Analysis (PCA) is used to reduce the dimensionality of the SME normalized matrix, and then non-linear UMAP embedding is used to generate the k-nearest neighbor (kNN) graph. kNN’s graph adjacency matrix is then clustered using Louvain clustering or k-means clustering. In the second stage, spatial information is utilized to identify sub-clusters from large clusters that span two or more physically distinct places. Using each location’s spatial coordinates, a two-dimensional k-d tree neighbour search is conducted.

## Cell Type Deconvolution

Cell-type deconvolution is the task of estimating cell-type composition in cell-pools from their aggregate transcriptomic information. This is a type of inverse problem, as we are trying to determine the signal of individual cell-types from aggregated readings across multiple cell-types. Moreover, due to the nature of the spatial (or bulk) transcriptomics profiling technologies, the true cell-type compositions are most often not given. Solving this task then requires some transfer learning approaches, in most cases using scRNA-seq data as a reference (transfer source).

The problem is formulated as follows. We’re given mixed-cell expression data *X* ∈ ℝ^*d*×*n*^ where each cell-pool *i* ∈ [1, *n*] is composed of a mixture of cell-types [1, *K*]. Then for each cell-pool *i* ∈ [1, *n*], we wish to construct an estimator *ŷ_i_* ∈ *Δ*^*K*−1^ of the true cell-type composition *y_i_* ∈ *Δ*^*K*−1^. Here, *Δ*^*K*−1^ is the regular *K*-simplex

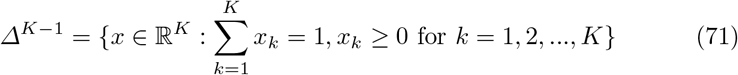

We are also given some reference scRNA-seq expression data *X*_s_ ∈ ℝ^*d*×*N*^ with one-hot labeled cell-types *C* ∈ ℝ^*N*×*K*^. Then construct the estimator of cell-type compositions for the *n* cell-pools by some function

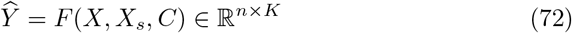

If the spatial information (2D or 3D coordinates in the given tissue)

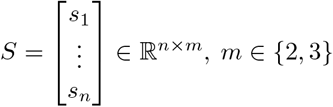

of the cell-pools are available, then we may incorporate this into the estimator of cell-type compositions for the *n* cell-pools by some function

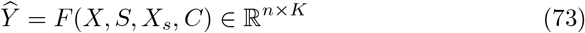

For the task of cell-type deconvolution, DANCE supports 4 models, one GNN based model and 3 non-GNN based models with classical regression models as their backbone.

## dance.modules.spatial.cell_type_deconvo.dstg

DSTG [120] is a GNN based method that constructs a graph from similarities between real mixed-cell expression data *X* ∈ ℝ^*d*×*n*^ and pseudo mixed-cell expression data 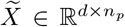 from reference scRNA-seq expression data *X_s_* ∈ ℝ^*d*×*N*^. First, the pseudo mixed-cell expression data is generated taking *n_p_* random samples (with replacement) of 2 to 8 cells from the scRNA-seq reference, and aggregating their UMI counts, downsampling to adjust for realistic bulk UMI counts. The pseudo and real mixed-cell data are then aligned in a lower dimensional (*S* < *d*) gene-space using Canonical Correlation Analysis (CCA). The projections to the *s* = 1, 2, … *S* dimensions are given by the canonical variables

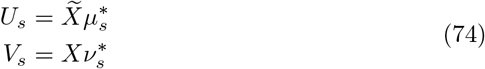

where

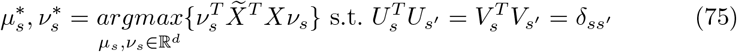

are the canonical correlation vector pairs. These embeddings are then used to construct a graph by considering Mutual Nearest Neighbors (MNN) as adjacent in the graph. That is, given a pair of sample cell-pools *i*, *j*, we let

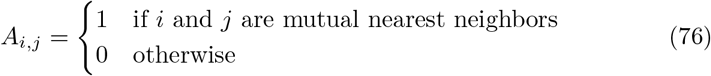

Here, adjacencies can be between simulated-to-real and real-to-real samples. With 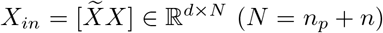 and the normalized adjacency matrix *Ã* as input, the *L* ≥ 1 (default 1) graph convolution (GCN) layers of the DSTG model are given by

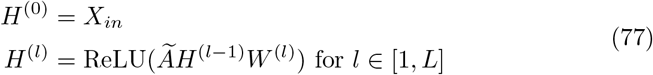

where *W*^(*l*)^ is the weight matrix for the *l_th_* layer. The output of the DSTG model is the predicted composition of *K* cell-types, given by

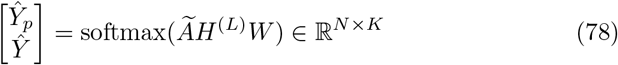

where *Ŷ_p_* and *Ŷ* are the predictions for the pseudo and real cell-pools, respectively. The loss function is then defined as the cross-entropy between the predicted and true cell-type compositions of the pseudo cell-pools

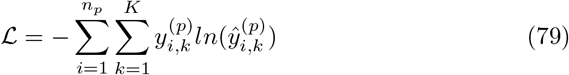

where 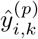 and 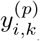 are the predicted and true composition of cell-type *k* in the *i_th_* pseudo cell-pool.

## dance.modules.spatial.cell_type_deconvo.spotlight

SPOTlight [38] builds on the classic non-negative least squares approach to cell-type deconvolution by incorporating topic-modeling for both the reference scRNA-seq expression data *X_s_* ∈ ℝ^*d*×*N*^ and the mixed-cell expression data *X* ∈ ℝ^*d*×*n*^. First, non-negative matrix factorization (NMF) is applied to the reference *X_s_* to get

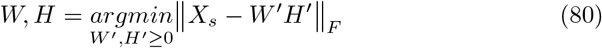

where the rows of *H* ∈ ℝ^*K*×*N*^ are the cell-topic embeddings, and the columns of *W* ∈ ℝ^*d*×*K*^ the corresponding weightings. Cell-topic profiles 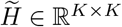 are then constructed from *H* by taking the median over each cell-type. Next, spot-topic profiles *P* ∈ ℝ^*K*×*n*^ are constructed through NNLS of *X* onto *W*

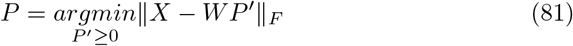

Finally, the estimator of cell-type compositions for the n cell-pools is then given by

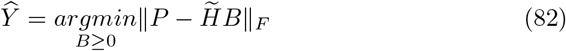

## dance.modules.spatial.cell_type_deconvo.spatialdecon

SpatialDecon [30] is non-negative linear regression based method that assumes a log-normal multiplicative error model between the mixed-cell data *X* ∈ ℝ^*d*×*n*^ and a cell-profile (signature) matrix 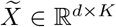. The cell-profile matrix 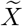 is a measure of center (median by default) expression for each of the *K* cell-types, constructed from the reference scRNA-seq data *X_s_* ∈ ℝ^*d*×*N*^. The log-normal multiplicative error model is given by

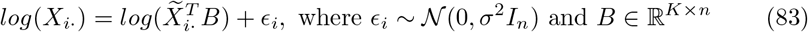

The estimator of cell-type compositions for the n cell-pools is then given by

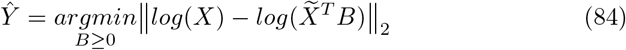

## dance.modules.spatial.cell_type_deconvo.card

CARD [85] applies a conditional autoregressive (CAR) assumption on the coefficients of the classical non-negative linear model between the mixed-cell expression *X* and a cell-profile matrix 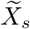, constructed from reference scRNA-seq *X_s_*. The linear model is given by

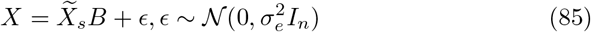

The CAR assumption then incorporates 2D spatial information 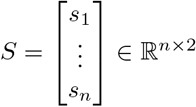 through an intrinsic prior on the cell-type compositions (the model coefficients) by modeling compositions in each location as a weighted combination of compositions in all other locations. This modeling assumption is given by

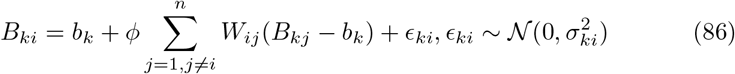

where the weights *W_ij_* are given by the Gaussian kernel

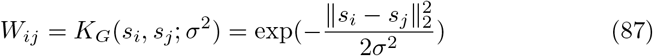

with default scaling parameter *σ*^2^ = 0.1. CARD then estimates the cell-type composition of the *n* cell-pools through constrained maximum likelihood estimation.

## E Structure Tree

The structure tree below shows our codebase structure in DANCE. Basically it consists of two parts: dance source code and examples. In example folder, each file represent one example to show how to leverage one specific model on dataset.

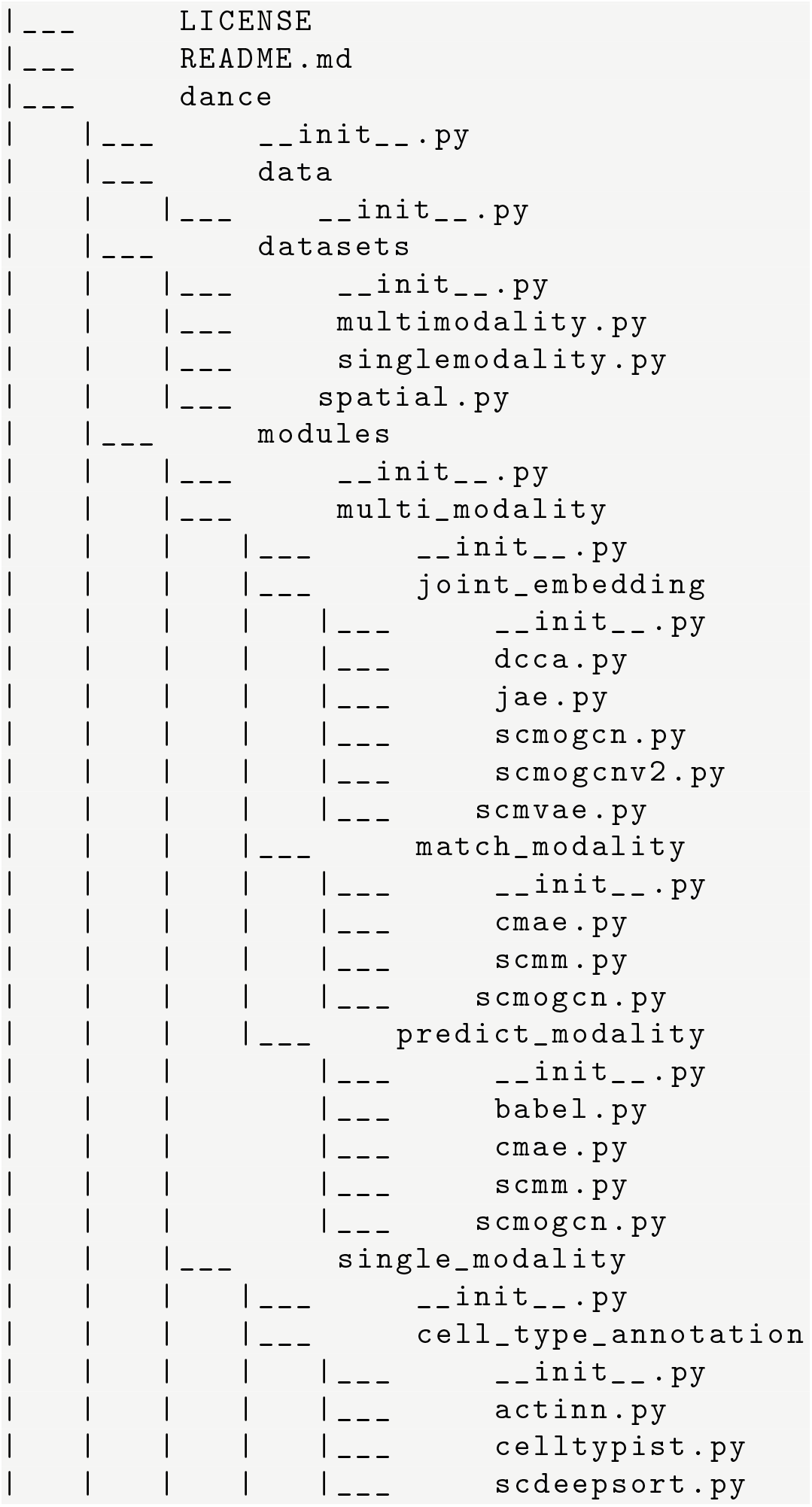

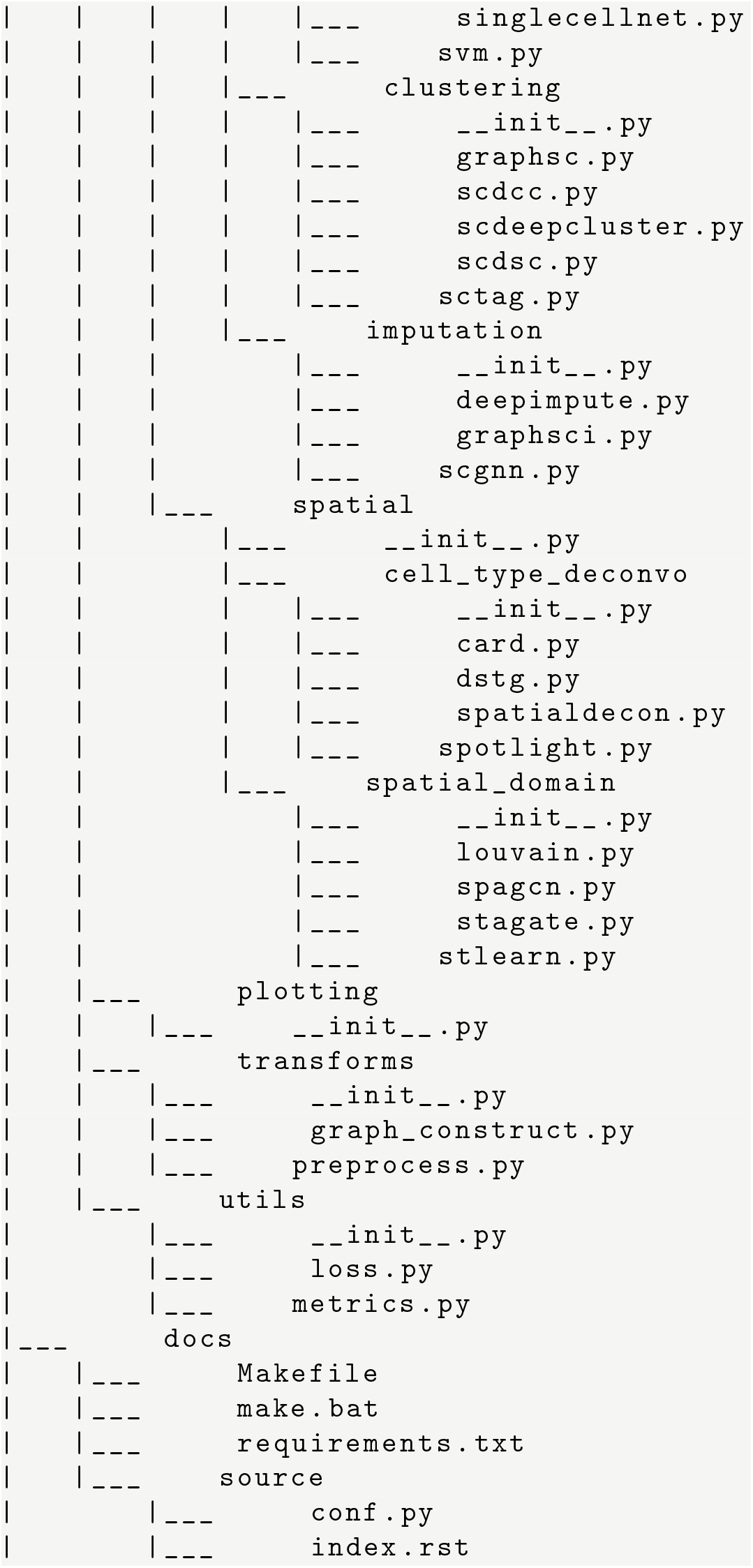

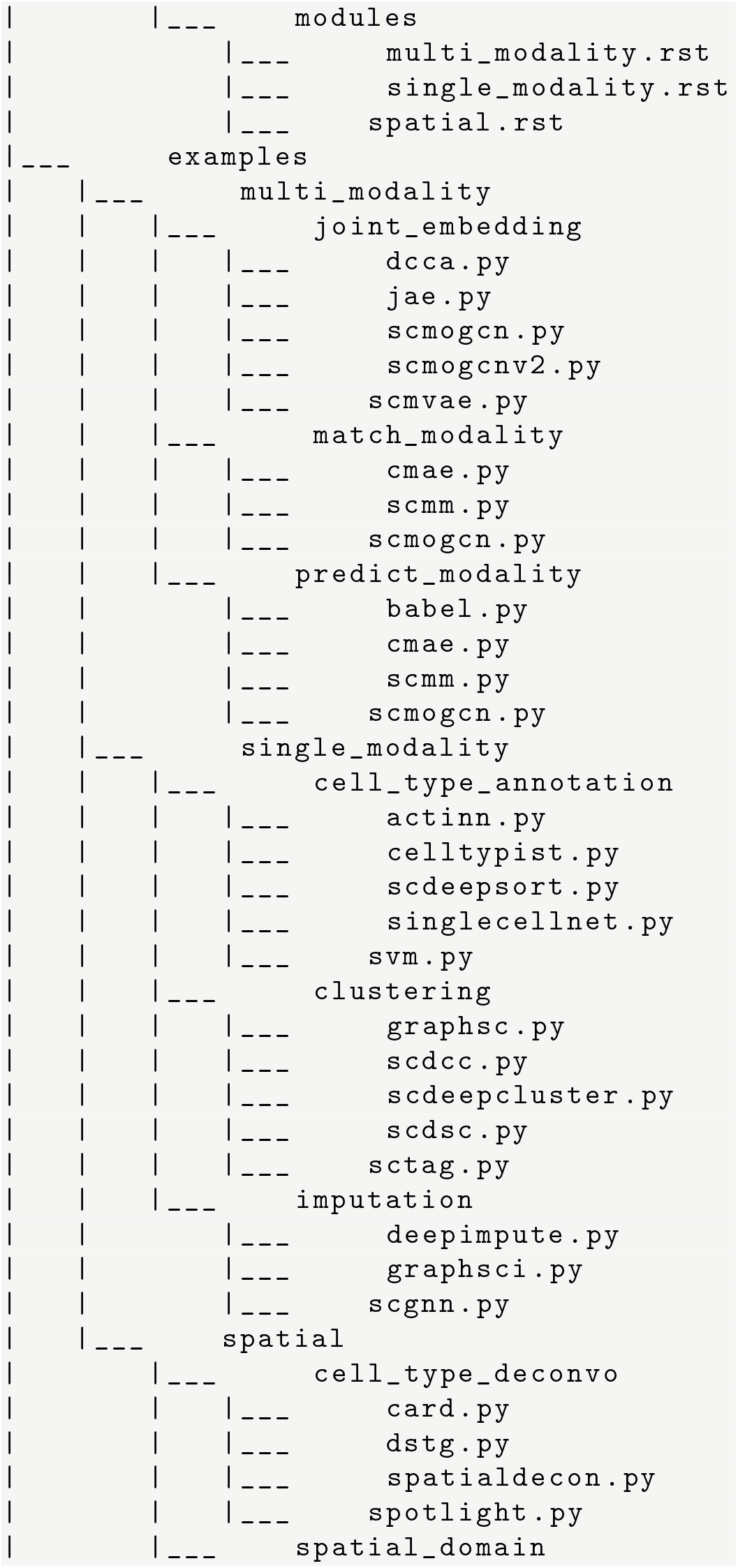

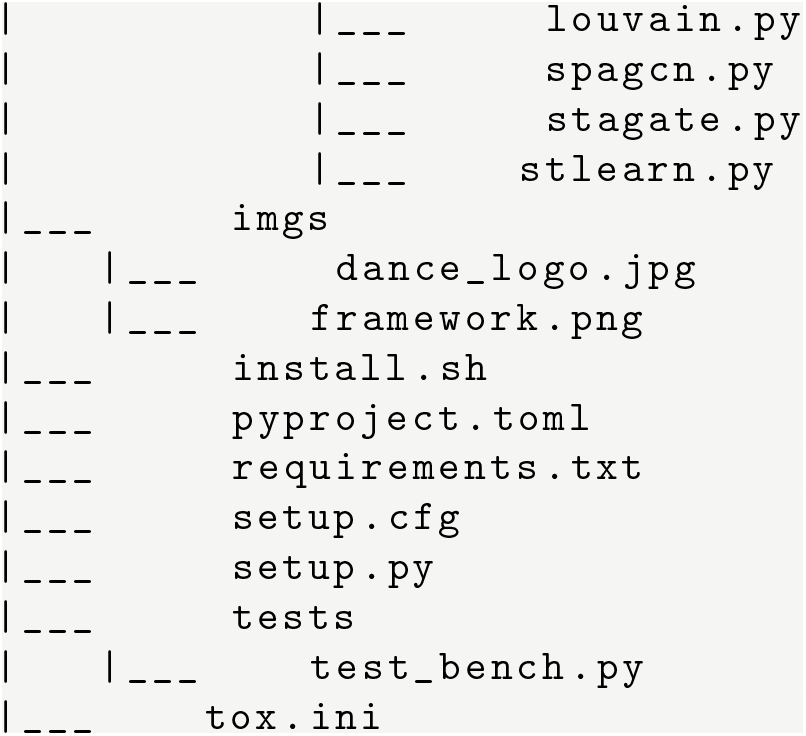

## Notes

### Competing Interest Statement

The authors have declared no competing interest.

### Summary of Updates

minor modification of the title

